# CoCoRV: a rare variant analysis framework using publicly available genotype summary counts to prioritize germline disease-predisposition genes

**DOI:** 10.1101/2021.09.29.462472

**Authors:** Wenan Chen, Shuoguo Wang, Saima Sultana Tithi, David Ellison, Gang Wu

**Affiliations:** Center for Applied Bioinformatics, St. Jude Children’s Research Hospital, Memphis, Tennessee, United States; Department of Computational Biology, St. Jude Children’s Research Hospital, Memphis, Tennessee, United States; Department of Cell & Molecular Biology, St. Jude Children’s Research Hospital, Memphis, Tennessee, United States; Department of Pathology, St. Jude Children’s Research Hospital, Memphis, Tennessee, United States

## Abstract

Sequencing cases without matched healthy controls hinders prioritization of germline disease-predisposition genes. To circumvent this problem, genotype summary counts from public data sets can serve as controls. However, systematic inflation and false positives can arise if confounding factors are not addressed. We propose a new framework, <u>co</u>nsistent summary <u>co</u>unts based <u>r</u>are <u>v</u>ariant burden test (CoCoRV), to address these challenges. CoCoRV has consistent variant quality control and filtering, ethnicity-stratified rare variant association test, accurate estimation of inflation factors, powerful FDR control, and can detect rare variants in high linkage disequilibrium. When we applied CoCoRV to pediatric cancer cohorts, the top genes identified were cancer-predisposition genes. We also applied CoCoRV to identify disease-predisposition genes in adult brain tumors and amyotrophic lateral sclerosis. Given that potential confounding factors were well controlled after applying the framework, CoCoRV provides a cost-effective solution to prioritizing disease-risk genes enriched with rare pathogenic variants.

## INTRODUCTION

To identify genetic variants, especially rare variants, that are associated with various human diseases, scientists have generated whole-exome sequencing (WES) and whole-genome sequencing (WGS) data, such as the Genome Aggregation Database (gnomAD)^1^, the Trans-Omics for Precision Medicine (TOPMed) program^2^, and the 100,000 Genomes Project^3^. Detecting causative rare variants typically requires a much larger sample size. Such a study should include at least thousands of cases and matched controls to ensure statistical power. However, most sequencing studies focused on a specific disease or trait include relatively few cases and very few (if any) matched healthy controls. Although many large cancer genomics studies have characterized the landscape of somatic mutations by sequencing germline and somatic samples from each patient, these studies do not include independent controls because germline samples from the same individuals are used as paired normal controls. Therefore, most cancer genomics sequencing studies cannot be used directly to perform germline-based case-control association analyses for discovery of novel cancer predisposition genes.

Combining the cases under scrutiny with external controls for prioritizing disease-predisposition genes is one solution. The best approach to controlling for confounding batch effects caused by the heterogeneity of exome-capture protocols, sequencing platforms, and bioinformatics-processing pipelines would be to re-map all raw sequencing data and then jointly call the genotypes of cases and controls. However, this approach is expensive and requires a lot of storage and a long processing time, which might not be feasible for some research groups. Another solution would be to download jointly called control genotype matrices (if available), merge with case genotype matrices, and apply variant quality control (QC) and filtering to account for batch effects. Alternatively, when the full genotype data are not available and assuming the pipelines have negligible effects on the association analyses after QC and filtering, one can use publicly available summary counts for case-control association analyses^4^. Here the summary counts refer to the allele or genotype counts of each variant within a cohort. If confounding factors are well controlled, the summary counts–based strategy is the least expensive and can dramatically increase the sample size of the controls. For instance, gnomAD v2.1 has more than 120,000 WES samples, and gnomAD v3 has more than 70,000 WGS samples^1^.

Using high-quality summary counts has led to important supporting evidence in identifying pathogenic germline mutations in adult cancers^5^ and pediatric cancers^6^. However, research is lacking on developing a general framework for using high-quality summary counts and evaluating the performance of such a framework.

Several challenges remain when using public summary counts for rare variant association tests. First, because the genotypes of the cases and controls are called separately, the QC and filtering steps might not be consistent if performed separately for cases and controls. To account for such differences, Guo *et al*.^4^ relied on the variant annotation QD (quality score normalized by allele depth); they also used synonymous variants to search for threshold combinations that would remove systematic inflation. One disadvantage of this approach is that it relies on one variant QC metric, instead of more advanced variant QC methods, such as VQSR^7^.

Differences in the genetic composition of unmatched cases and controls present another challenge. When using publicly available summary counts, researchers often use Fisher’s exact test (FET)^4,6^ which either treats different ethnicities as a single population or selects samples from one major ethnicity, e.g., samples of European ancestry. Ignoring ethnicity when using public summary counts can result in false positives due to population structure when rare variants have relatively high minor allele frequencies (MAFs) in a specific population. On the other hand, considering a single matched ethnicity might reduce the statistical power. Although a better, more common practice is to use all samples and account for the population structure by adjusting for the principal components (PCs)^8^, it is not applicable when only public summary counts are available without individual genotype information.

The inflation factor estimation of the association test when focusing on very rare variants is also important. In genome-wide association studies with common variants, the null distribution of p-values is assumed to be continuous and follow a uniform distribution *U*(0, 1). However, for rare variants, especially those with very low MAFs, assuming a uniform distribution of the p-values under the null hypothesis of no association is no longer accurate. For example, in the FET under the null hypothesis of no association, the count of cases with alternate alleles in a 2×2 contingency table for each gene follows heterogenous hypergeometric distributions, depending on the total number of rare alleles. For the same reason, traditional FDR control method such as the BH procedure^9^ assuming a uniform distribution of the p-values under the null is suboptimal for the discrete count based test results^10^.

Lastly, the most critical challenge when using public summary counts is related to rare variants in high linkage disequilibrium (LD). In most cases, the independence assumption is probably reasonable because the variants are rare and the chance of observing both variants in one sample is low, unless they are in strong LD. However, when the assumption is violated, either due to high LD or technical artifacts, that can cause false-positive findings or systematic inflation. Here we propose a framework that better prioritizes potential causal genes by using publicly available summary counts. We focus on genotype calls from WES data throughout this study.

## RESULTS

### Overview of the proposed framework

To address the challenges in rare variant association tests using public summary counts, we developed a new framework termed <u>Co</u>nsistent summary <u>Co</u>unts based <u>R</u>are <u>V</u>ariant burden test (CoCoRV) (Fig. 1). The input case data can be either full genotype based or summary count based. The input control data are summary count data such as from gnomAD. We included a fast and scalable tool to calculate the coverage summary statistics because coverage depth is important in consistent filtering between cases and control^4^. Cases belonging to different ethnicities can be matched to corresponding controls. CoCoRV allows a user to customize criteria for filtering variants based on annotations in the input data. We provided several built-in variant categories based on annotated functions from ANNOVAR^11^ and REVEL scores^12^. For putative pathogenic variants, we included variants annotated with “stopgain”, “frameshift_insertion”, “frameshift_deletion”, and “nonsynonymous” with a REVEL score ≥0.65, which showed empirically good discrimination power for potential pathogenic versus nonpathogenic variants.

**Fig. 1.**
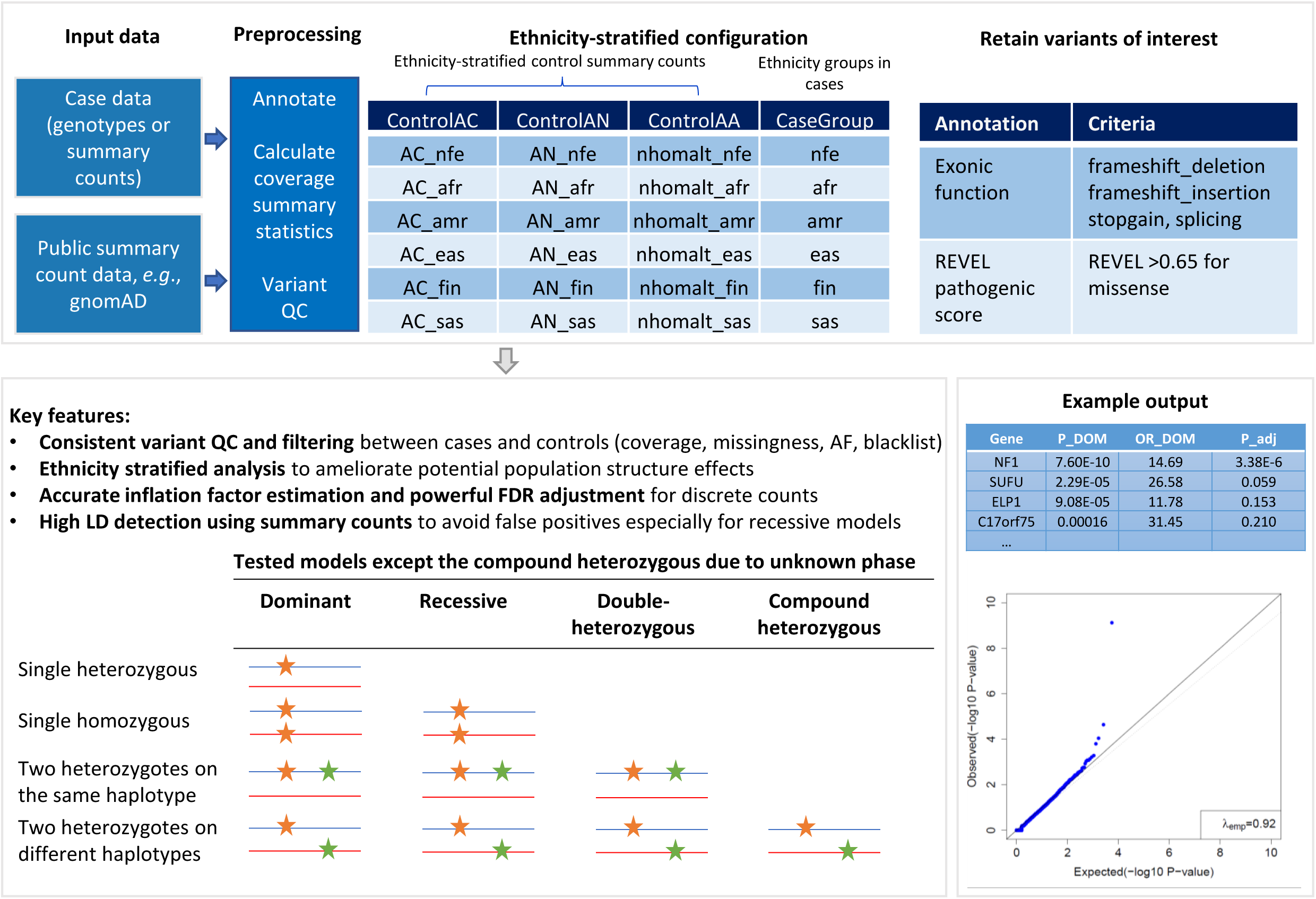
Overview of the proposed framework CoCoRV. The top box describes the main input and preprocessing. The bottom left box describes the key features and the tested models. The bottom right box shows the main output. ControlAC: the annotation for the alternate allele count; ControlAN: the annotation for the total allele count; ControlAA: the annotation for the homozygous alternate genotype count; P_DOM: p-value of the dominant model, OR_DOM: the odds ratio of the dominant model; P_adj: the adjusted p-values for multiple testing.

CoCoRV ensures that the variant QC and filtering are consistently applied between cases and controls. In addition to the FET, CoCoRV includes ethnicity-stratified analysis using the Cochran–Mantel–Haenszel (CMH)-exact test, which mitigates systematic inflations when samples include multiple ethnicities. We computed sample counts in three models (dominant, recessive, and double-heterozygous) within each gene.

Due to the discrete nature of count data, we propose an accurate inflation factor estimation method that is based on the sampling the true null distribution of the test statistics (Fig. 2a) and is essential for checking possible systematic inflations. Similarly, the commonly used BH procedure^9^ for FDR control assumes a continuous uniform distribution, which is not true for discrete count based tests. We therefore propose to use more powerful resampling based FDR control methods for the FET and CMH test results (Fig. 2a). The sampled p-values under the null hypothesis can be used for both inflation factor estimation and resampling based FDR control.

**Fig. 2.**
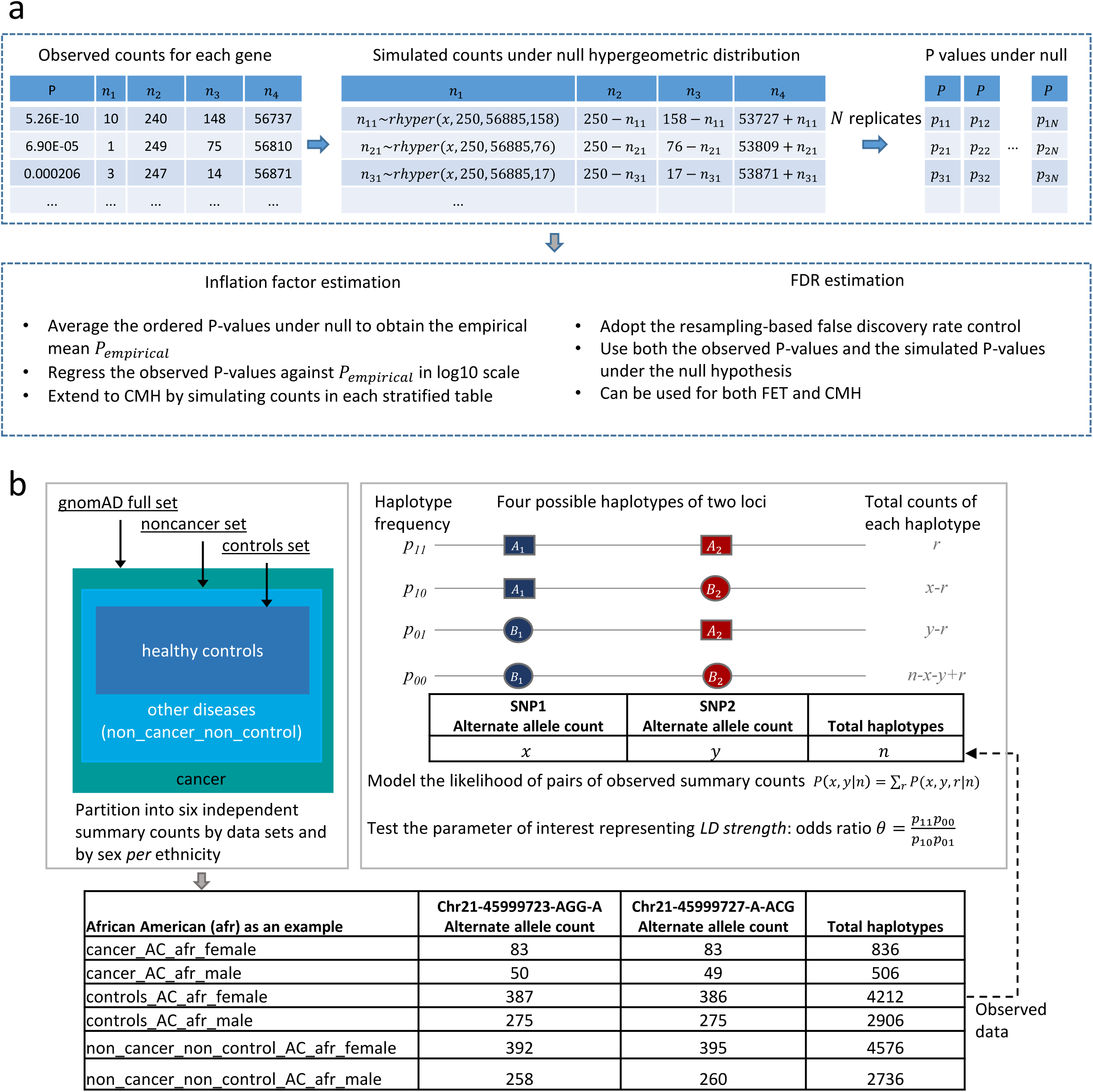
The schematic representation of proposed methods in CoCoRV. a. The estimation of inflation factors and FDR. The sampled p-values under the null can be used for both inflation factor estimation and FDR control. b. LD detection using gnomAD summary counts. The gnomAD full data sets were partitioned into three independent sets of summary counts and then further partitioned into six independent sets stratified by sex. The six independent summary counts between a pair of variants can be modeled with the LD strength as a parameter, and the LD strength can be tested for LD detection. The table at the bottom shows an example of the six independent summary counts between two variants of high LD in the African American ethnicity group.

We introduce a LD-detection method using only gnomAD summary counts to identify high-LD variants (Fig. 2b). It partitions the gnomAD data set into several independent summary count sets, and then infers the high-LD between variants based on the generated independent summary counts for each ethnicity. Excluding high-LD variants results in a more accurate estimation of the number of samples in each model and reduces false positives when the assumption of independence between variants is violated, especially for the recessive and double-heterozygous models. Details of the framework and modeling are provided in Methods.

### CoCoRV’s inflation factor estimation is unbiased and accurate

We compared our proposed inflation estimation method with two other methods from TRAPD^4^. We used the rare variant analysis results of the central nervous system tumors (CNS) and acute lymphoblastic leukemia (ALL) cohorts against our in-house controls under a dominant model to compare these methods (Fig. 3). We also checked their distributions under the simulated null p-values, where a well-estimated inflation factor should be unbiased and centered around 1. For the CNS cohort tested using FET (Fig. 3a), the inflation factor *λ*_*emp*_ estimated using empirical null p-values was 0.96, and those of the other two methods were greater than 1. For the ALL cohort tested using FET (Fig. 3b), all three inflation factors were less than 1. The rightmost plots in Fig. 3 show that *λ*_*emp*_ was well calibrated under the null hypothesis, while the other two methods from TRAPD showed either upward or downward biases. The QQ plots based on CoCoRV’s method showed almost straight lines along the diagonal, and the other two methods showed clear shifts from the diagonal. In addition to FET, we saw similar patterns using the CMH-exact test (Fig. 3c, d). The estimation of CoCoRV’s *λ*_*emp*_ was stable, even with as few as 100 simulated null p-values. Therefore, in later estimations of *λ*_*emp*_ in QQ plots, we used 100 simulated null p-values *per* gene, unless otherwise specified. When the null distribution of the inflation factor is of interest, as shown in the rightmost plots, it is better to use at least 1,000 replications for a more stable result. Because of the better performance of the proposed estimation, we applied it to nearly all subsequent analysis results in this study. The run time of our inflation factor estimation depended on the number of variants, their AFs, and the number of genes tested. In this study, the time to run 100 replications was several minutes using one CPU core, and there is an option to run the estimation on multiple cores to further speed up the process.

**Fig. 3.**
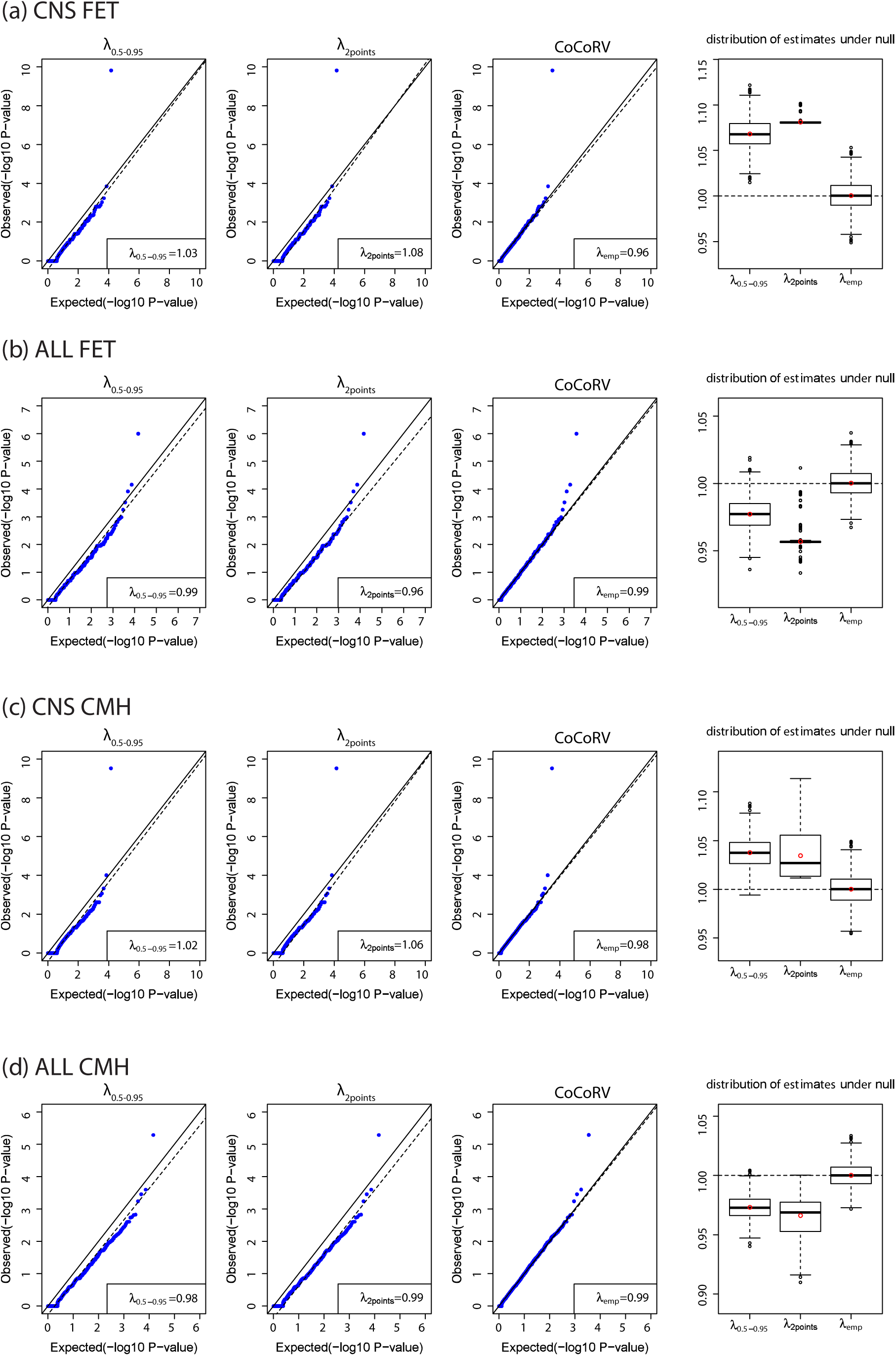
Comparisons of different inflation factor estimation methods. The first three columns show the inflation factors estimated using different methods. The dashed lines are the fitted lines in the inflation factor estimation. The fourth column shows boxplots of the estimated inflation factors using simulated p-values under the null hypothesis. The red circle indicates the mean value, a. Results based on the central nervous system (CNS) cancer cohort and Fisher’s exact test (FET). b. Results based on the acute lymphoblastic leukemia (ALL) cohort and FET. c. Results based on the CNS cohort and the Cochran−Mantel−Haenszel exact test (CMH) stratified by ethnicities, d. Results based on the ALL cohort and the CMH stratified by ethnicities.

### Concordance of rare variant association between using summary counts and full genotypes

We compared the concordance between the result obtained using summary counts from separately called controls and that using jointly called full genotypes. For this analysis, we generated separately called control summary counts using an in-house control cohort of 8,175 WES and treated the jointly called case-control full genotypes as the ground truth (Fig. S1). To our knowledge, this is the first direct comparison between analysis results using summary counts of separately called cases and controls and that using jointly called full genotypes. For this comparison, we focused on the commonly used dominant model. We used an AF threshold of 1*e*–3 for the CNS and ALL cohorts and for all comparisons.

We first compared the sample counts with defined pathogenic variants for each gene. For cases, we compared and tabulated the sample counts with defined potential pathogenic variants from using jointly called full genotypes and that from using summary counts into a cross-classification/contingency table (Fig. 4a, b). For both the CNS and ALL cohorts, the concordance between using full genotypes and CoCoRV was very high (Pearson’s correlation r >0.98). Most of the discordances were within one count. For controls, because the number of samples with defined potential pathogenic variants varied in a large range, we used the scatter plot of those numbers to illustrate the concordance between using full genotypes and using summary counts (Fig. 4c, d). The correlation between using full genotypes and CoCoRV was also very high (r = 0.996). The high correlation of the raw counts shows that the proposed summary counts–based framework maintained a high concordance with the jointly called full genotype–based framework. We also examined the top genes of the association tests between using jointly called full genotypes and separately called summary counts (Fig. 4e, f). In general, the percentage of top overlapped genes was about 70% or higher, especially for the more stringent p-value thresholds. We also performed ethnicity-stratified analyses for both frameworks and found similar overlapping patterns (Fig. S2). Finally, the QQ plots using jointly called full genotypes and separately called summary counts were also similar, with correlations between –log_10_ p-values greater than 0.9 (Fig. 4 g, h).

**Fig. 4.**
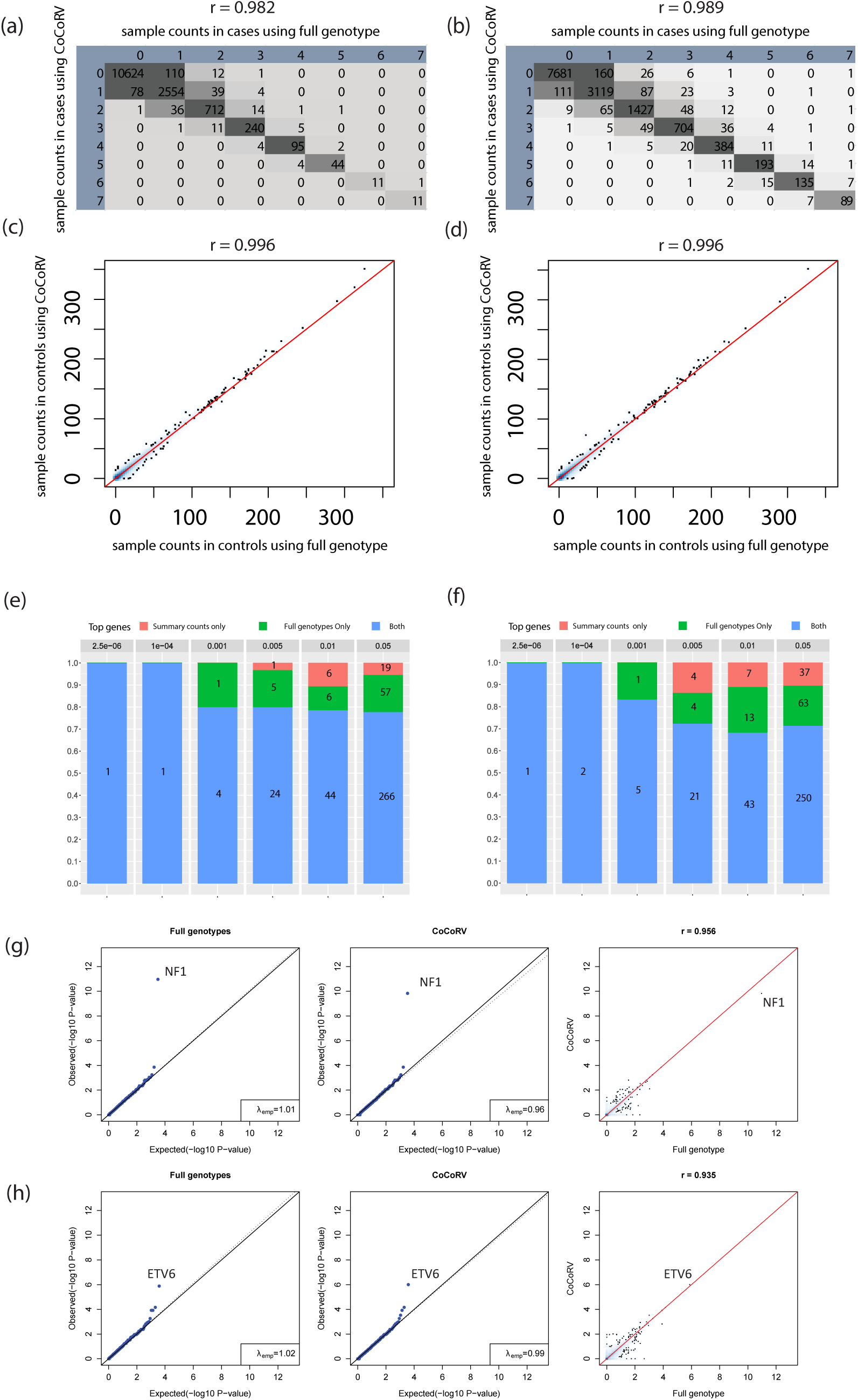
Comparisons of concordance between using separately called summary counts and jointly called case-control full genotypes. a-b. Cross-classification tables between using full genotypes and using summary counts across all genes on the number of samples carrying defined pathogenic alleles in CNS (a) and ALL (b) cohorts. The column ID is the number of samples calculated using full genotypes, and the row ID using summary counts. c-d. Scatter plots of the number of control samples from CNS (c) and ALL (d) cohorts carrying qualified rare alleles, by using full genotypes and summary counts. e-f. Comparisons of top genes in the CNS (e) and ALL (f). The bar heights show the percentage, and the numbers show the absolute number of genes. g-h. QQ plots of association results using full genotypes (left panels), summary counts (middle panels), and scatter plots of-log 10 p-values between using full genotypes and summary counts. Results are from CNS (g) and ALL (h).

### FDR methods accounting for discrete counts show good FDR control and improved power

Two adopted resampling based methods: RBH_P and RBH_UL, and one recently developed method ADBH.sd^10^ show good FDR control in general and similar improved power for FET based test results, even though RBH_P has slightly inflated FDR in some settings (Fig. S3). Compared to BH, the relative power increases of RBH_UL range from 11% to 27% (Table S1). Filtering the genes with few rare allele counts (BH_T2 and BH_T3) shows little power increase. When applied to the CNS association test results using gnomAD as the control (threshold <0.2), the three methods accounting for the discrete counts all identify the established genes *NF1, SUFU, ELP1* as significant, while the BH method only identifies *NF1* as significant (Table S2). The resampling methods we adopted are more general and can be applied to both the FET and the CMH test results, while ADBH.sd supports FET but not CMH test results. The time cost is similar as that in inflation estimation because sampling the p-values under the null hypothesis is main time consuming part.

### The proposed LD detection method has high power in detecting high-LD rare variants and accurately identifies MNVs in gnomAD

We first evaluated the type I error and statistical power of the proposed LD test by using simulated data consisting of six independent sets of summary counts (Fig. S4a). The type I error was well controlled. The power of detecting LD increased as the LD strength and the AF increased. The LD test was well powered for detecting strong positive LDs but not for detecting weak LDs. When full genotypes are available, the proposed test can also be used (Fig. S4b). Simulations based on full genotypes showed that the type I error was also well controlled. The power increased remarkably compared to using summary counts, which was expected given that more information is available when using full genotypes.

As one application and a validation of the proposed LD detection method, CoCoRV was applied to detect high-LD multi-nucleotide variants (MNVs)^13^ without access to the sequencing reads information. A MNV is defined as a cluster of two or more nearby variants on the same haplotype in an individual. Table 1 summarizes the relation between detecting high-LD variants and MNVs. We applied our proposed LD test to scan the whole exomes of gnomAD data to detect rare variant pairs of high LD and compare them with the reported gnomAD MNVs detected using sequence reads. Although high-LD variants and MNVs are not the same, they have similarities when restricted to rare variants with distances ≤2 BPs (Table 1). For each ethnicity and each variant, we extracted the gnomAD exome count data and generated six independent sets of summary counts using the cohort and sex information. Then we applied the LD test on pairs of variants annotated with certain functions, excluding synonymous variants.

**Table 1.** Relations between the detection of LD and that of gnomAD MNV

| Type of detection | LD | Distance | Input |
| --- | --- | --- | --- |
| LD | High, can be estimated in the test | No restriction | Summary counts or full genotype matrix of a cohort |
| MNV | Most high, but can be low or none, need to validate in a cohort | Small, often $\leq 10$ BPs | Sequencing reads from each individual bam file |

Abbreviations: BPs, base pairs; LD, linkage disequilibrium; MNV, multi-nucleotide variant

The detection results in gnomAD *per* ethnicity group are summarized in Table 2. In total, approximately 10 million tests of variant pairs were performed. By controlling the FDR at 0.05, we detected nearly 10,000 coding variant pairs that are in high LD (Table S3). Most of the detected variant pairs with FDR <0.05 were high-LD variants (odds ratio >1*e*5). We also checked the overlap between high-LD variants and reported gnomAD MNVs. Because the reported gnomAD coding MNVs are pairs of SNVs within 2 BPs, we restricted the detected pairs to high-LD (odds ratio >1*e*5) SNVs for both variants and the distance within 2 BPs. About 90% of them are reported in the gnomAD MNV data set (either from exomes or genomes) using read-based MNV detection^13^. We then manually checked the read haplotype information from the gnomAD website (https://gnomad.broadinstitute.org/) for those 115 pairs not reported as gnomAD MNVs. There were 83 unique pairs if not distinguishing ethnicities, and 65 (78.3%) had read information. Of the 65 pairs, 60 (92.3%) had supported reads showing that they are on the same haplotype. The reason that these pairs were not reported as gnomAD MNVs might be due to other filtering criteria used. Reported gnomAD MNVs had alternate alleles verified on the same haplotype, which provides strong evidence that the pairs detected through our LD test are true high-LD variants because the likelihood of two independent rare alleles appearing on the same haplotype is very low. Fig. S5 shows the distribution of distances and corresponding recombination rates between a pair of high-LD SNVs. Although there was a peak when the distance was <10 BPs, there was also a large mass when it was >10 BPs, emphasizing the importance of detecting high LD with relatively long distance. In general, the recombination rate decreased as the physical distance increased, indicating that even though the physical distance was large, the recombination frequency between the two variants can still be low. Over relatively long distances (e.g., >150 BPs), high-LD variants could not be detected by examining the sequencing reads data (usually ∼150 BPs) but could be detected using CoCoRV.

**Table 2.** LD detection of annotated functional variants within each gene by using gnomAD summary counts

| **Ethnicity** | **Sample size** | **Pairs tested** | **FDR <0.05** | **High LD<sup>a</sup>** | **MNV (≤2 BPs)** | **Reported in gnomAD MNV<sup>b</sup>** |
| --- | --- | --- | --- | --- | --- | --- |
| nfe | 56,885 | 4,760,061 | 1,958 | 1,912 | 311 | 278 (0.89) |
| afr | 8,128 | 1,166,251 | 3,788 | 3,525 | 284 | 264 (0.93) |
| amr | 17,296 | 2,171,082 | 1,945 | 1,925 | 200 | 181 (0.91) |
| eas | 9,197 | 483,555 | 922 | 878 | 135 | 118 (0.87) |
| sas | 15,308 | 1,407,370 | 970 | 953 | 119 | 97 (0.82) |
| fin | 10,824 | 177,390 | 510 | 488 | 79 | 75 (0.95) |

<sup>a</sup>High LD was defined as estimated odds ratio >1e5.
<sup>b</sup>The union of gnomAD coding MNV and genome MNV with distance ≤2 BPs. The numbers within the parentheses are the proportions of our detected MNV (≤2 BPs) reported in the gnomAD MNV data set.
Abbreviations: nfe, non-Finnish European; afr, African American; amr, Admixed American; eas, East Asian; sas, South Asian; fin, Finnish; BPs, base pairs; FDR, false discovery rate; MNV, multi-nucleotide variant.

The above analyses used the stringent threshold FDR <0.05 to achieve high precision but could sacrifice the recall rate. We also used different p-value thresholds and evaluated the recall and precision by treating the reported gnomAD MNVs (≤2 BPs) as the ground truth, though some high-LD MNVs detected were not reported in the gnomAD MNVs, as described above. In total, there were 1,780 reported gnomAD MNVs among our tested variant pairs (Table S4). If we chose 0.01 as the p-value threshold, 1,635 of 1,976 detected were true, corresponding to precision 0.83 and recall 0.92. The LD detection results, including p-values and FDRs, were stored in a file and used later in CoCoRV to exclude high-LD variants in controls and cases.

### Detecting high-LD variants improves count estimation and removes false positives in the recessive and double-heterozygous models

We performed a simulation of count estimation by using summary counts from a set of variants, including independent and correlated variants (Fig. S6). Our results showed that the count estimation results from using CoCoRV were more accurate than those from using TRAPD. TRAPD overestimated the counts for all three models when high-LD variants were present, most likely because TRAPD was designed to be conservative for a one-sided FET to mitigate the violation of the assumption of independent variants. This overestimation could result in an inflated type I error, and the potential loss of power if a two-sided test is used. CoCoRV corrected for high-LD variants and resulted in a much better estimation.

We also compared the association results under the recessive model, with or without the LD test with an AF threshold 0.01 and using gnomAD summary counts as controls for both the CNS and ALL cohorts (Fig. 5). When there was no LD check for cases with two heterozygous variants, we found several false positives with very low p-values (Fig. 5a, c). When variants in LD were excluded from the count of case samples (Fig. 5b, d), those false positives were removed. Applying the LD test also removed false positives with very low p-values under the double-heterozygous model (Fig. S7). We noticed that the QQ plots under the recessive model were not as well calibrated as those under the dominant model, even after applying the LD test. The reason for this result could be that not all double-heterozygous variants with positive LD were detected using gnomAD summary counts, as indicated in the power simulations. Another factor to consider is that the recessive and double-heterozygous models are sensitive to differences in LD structures between cases and controls. Stratifying samples into different ethnicities might still retain some LD differences due to the population substructure within each of the six ethnicities. For example, African/afr and Latino/amr are two main admixed populations, and different proportions of ancestries can change the LD structure. Although there is no perfect solution, our CoCoRV method represents an advance in using only summary counts as controls for rare variant association tests.

**Fig. 5.**
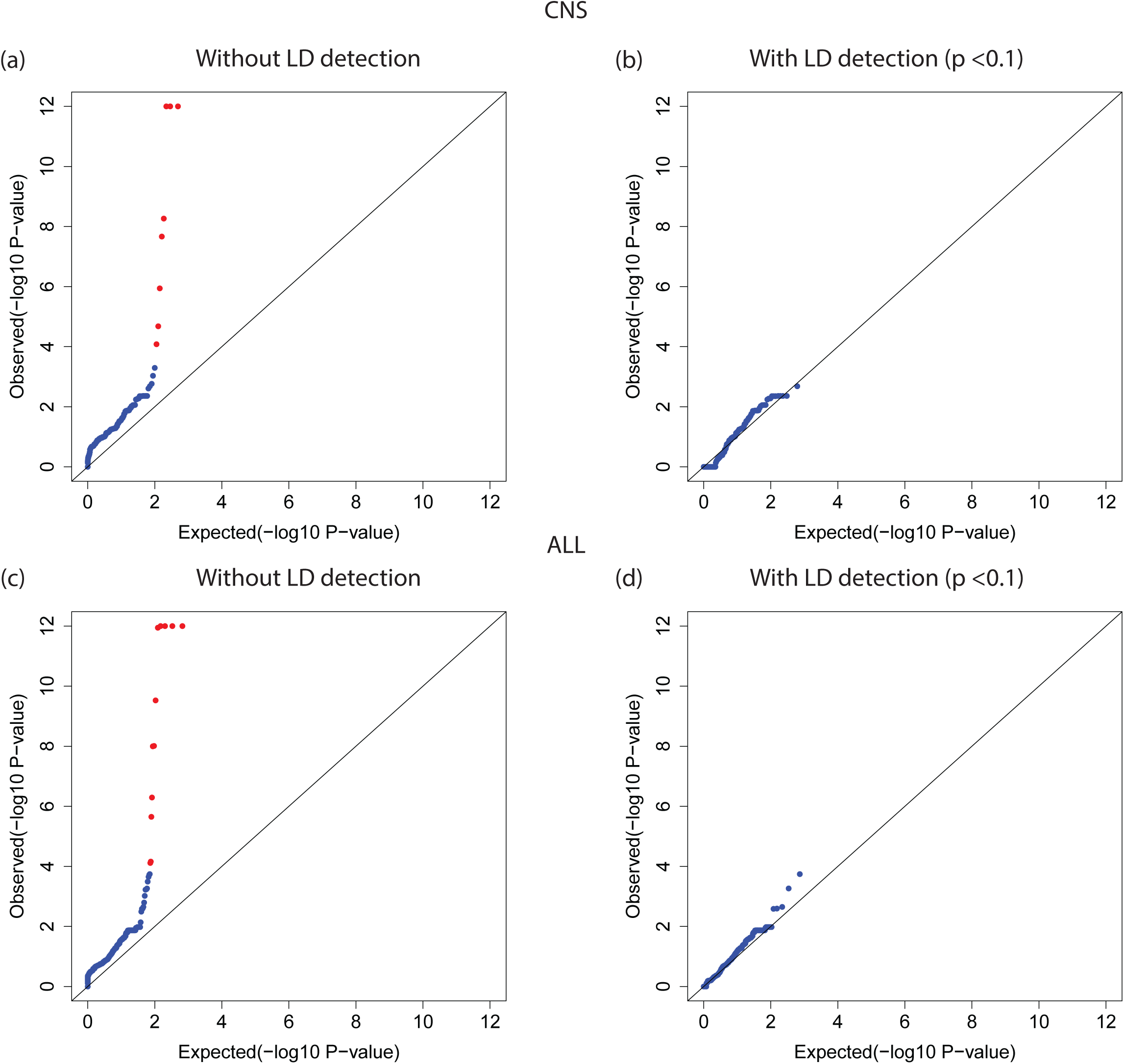
Employing LD detection removes false positives under the recessive model. a, c. QQ plots of association tests under the recessive model without employing LD detection in the CNS and ALL cohorts, respectively. b, d. QQ plots of association tests under the recessive model after employing LD detection with a p-value threshold of 0.1. Red dots in (a) and (c) are false positives that disappeared after employing the LD detection.

### CoCoRV analysis of the CNS and ALL cohorts

We applied CoCoRV to two St. Jude pediatric cancer cohorts (CNS and ALL) and used gnomAD summary counts as controls under a dominant model. For a comparison, we also applied TRAPD to the two cohorts. We used the all-pooled *AC* and *AN* to generate the counts per gene because TRAPD is hard-coded to use the annotation *AC* and *AN* to generate case-control statistics. CoCoRV was flexible to specify any annotated fields, such as non-cancer subsets, or any population-specific counts from gnomAD. Both SNVs and short indels were considered. For the CNS cohort, TRAPD showed an inflated QQ plot when using its own inflation estimator *λ*_*2points*_(Fig. 6a). After manual variant checking of the top six genes from TRAPD’s results, there were two false positives. These were mainly due to indels of bad quality that passed the QD threshold based on TRAPD’s suggested search strategy. VQSR correctly identified these variants as low-quality variants, and gnomAD’s random forest filter also flagged them as low-quality. This suggests that a single QD threshold is not sufficient, especially for indels in variant QC. CoCoRV showed no inflation in either the pooled counts from all ethnicities (Fig. 6b) or the stratified analysis using CMH (Fig. 6c), and it identified the known causal gene *NF1* as the top gene. For the ALL cohort, five false positives were among the top six TRAPD-discovered genes that were manually checked; all had problematic indels with similar reasons as those detected in the CNS cohort. There are also inflations in the QQ plot (Fig. 6d). In contrast, CoCoRV showed no obvious systematic inflation and identified the known causal gene *ETV6* as the top gene. The CMH-based analysis had better calibration at the tail than did the FET-based analysis (Fig. 6e, f).

**Fig. 6.**
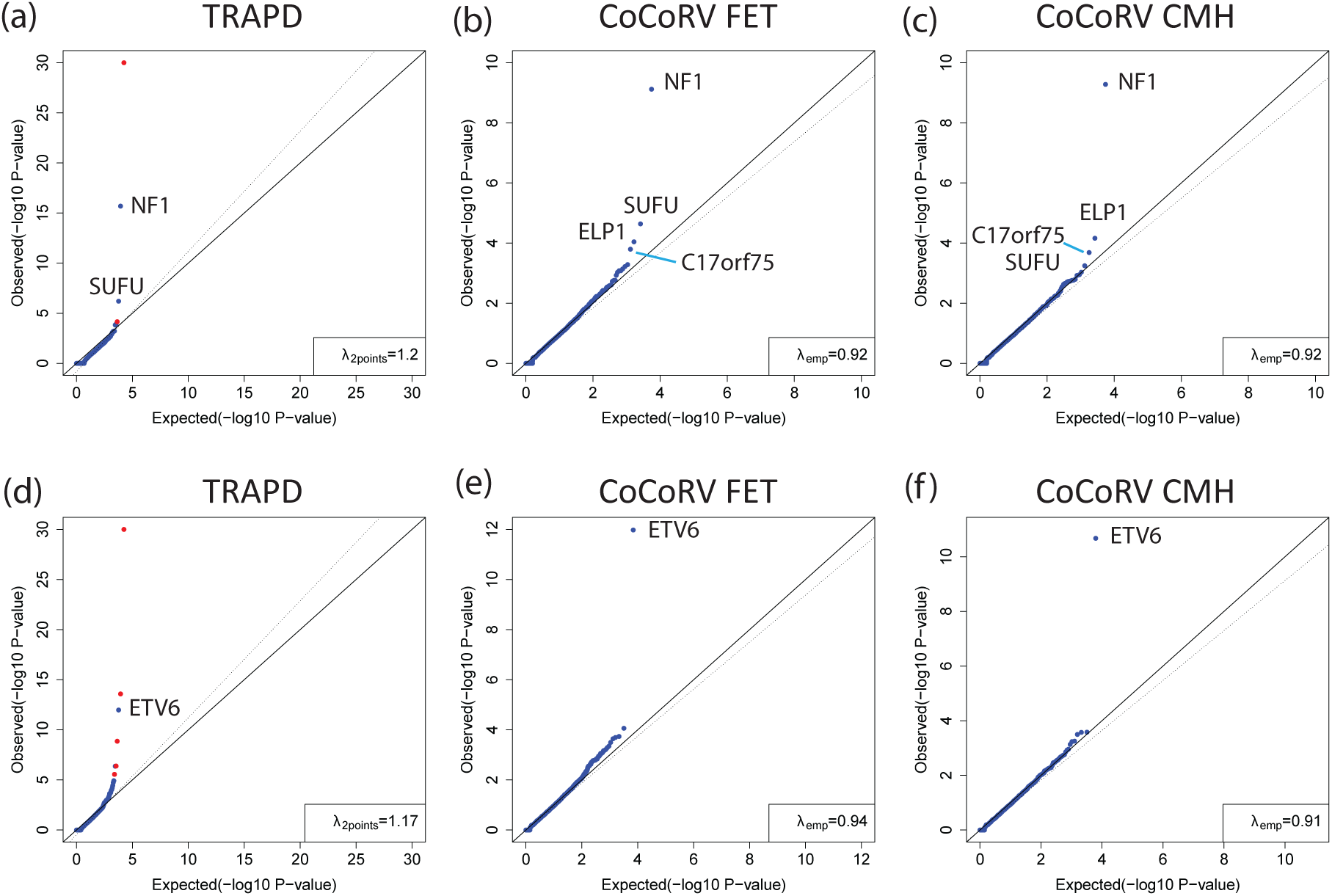
CoCoRV and TRAPD analyses of the CNS and ALL cohorts. We used gnomAD summary counts as controls. a-c. The analyses of the CNS cohort using TRAPD (a), CoCoRV with the Fisher’s exact test (FET) (b), and CoCoRV with the Cochran−Mantel−Haenszel (CMH) exact test (c). d-f. The analysis of the ALL cohort using TRAPD (d), CoCoRV with the FET (e), and CoCoRV with the CMH exact test (f). Red dots in (a) and (d) indicate genes that are false positives after manual check of the variants among the top six ranked genes.

*NF1* was detected in the pediatric CNS cohort and had relatively similar p-values across different scenarios, no matter which summary counts–based controls were used. For the ALL cohort, the much larger sample size of gnomAD dramatically increased the significance of *ETV6*, from 10^−6^ to 10^−12^ (Table 3). The relatively stable p-values for *NF1* in the CNS cohort were most likely due to the adequately large counts in the contingency table using the in-house– constructed summary counts. The benefit of increasing the number of controls also diminished after a certain level, while the number of cases was fixed (Table 3). In contrast, *ETV6* had 0 or 1 qualified control sample with rare alleles in the in-house controls, had more significant p-values by using gnomAD as controls, benefited from the improved quantification of the AFs when the control sample size increased substantially.

**Table 3.**
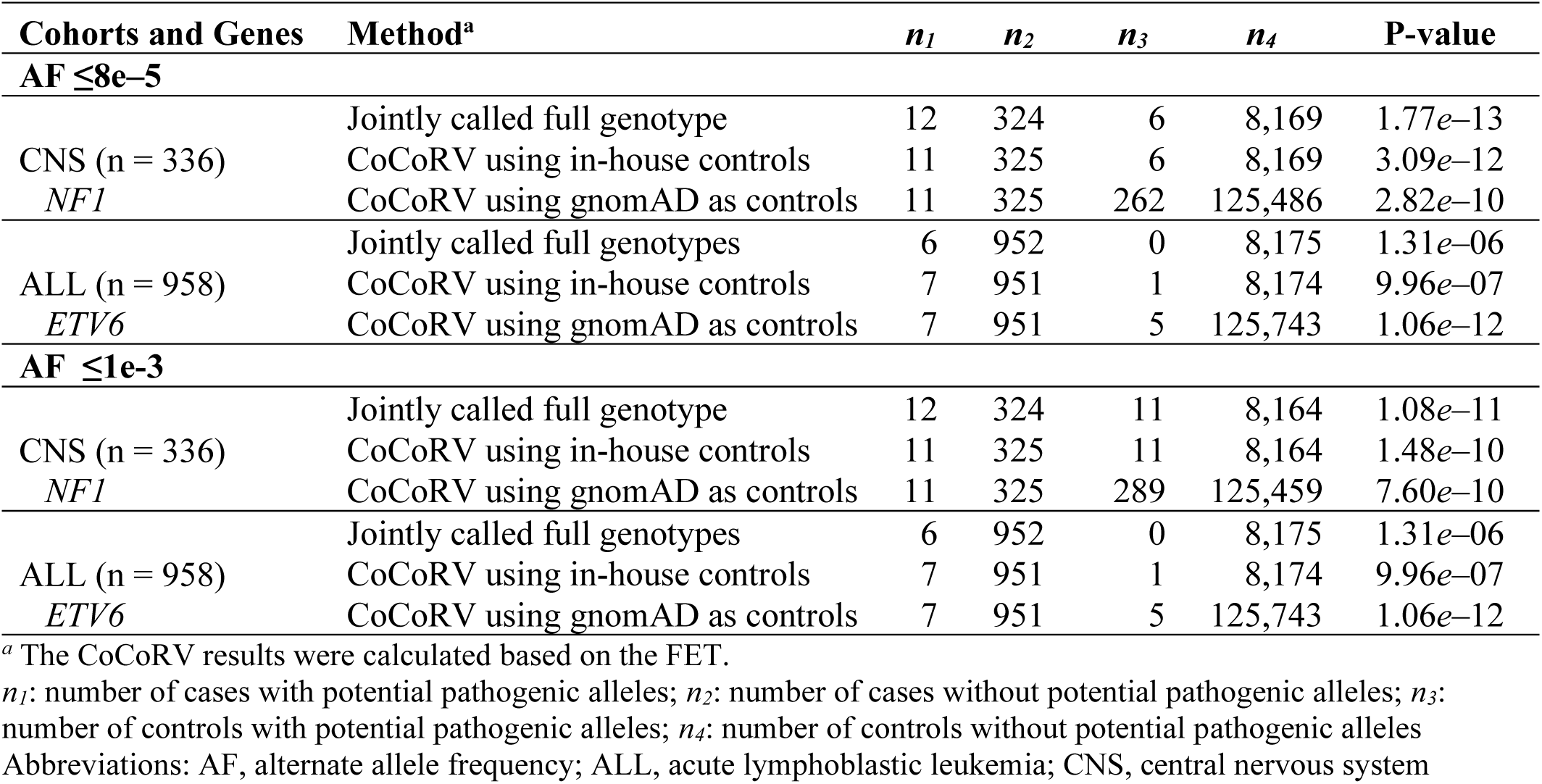
Statistics of the top genes by using different methods and alternate allele frequency thresholds

For the CNS cohort, the top four genes were *NF1, ELP1, C17orf75*, and *SUFU* (Fig. 6). *NF1* and *SUFU* are well established causative genes for pediatric brain tumors^14^. *ELP1* was recently identified as a predisposition gene of pediatric brain tumors^6^. This makes *C17orf75* an interesting candidate. From the GTEx portal, two of the top three tissues with highest median expression of *C17orf75* are the cerebral hemispheres and the cerebellum (Fig. S8). Further validation is needed to confirm the association of *C17orf75* with pediatric brain tumors.

### Analysis of germline samples from The Cancer Genome Atlas (TCGA)

We applied CoCoRV to the TCGA WES data sets of glioblastoma multiforme (GBM) and low-grade glioma (LGG). Brain tumors can be classified into grades I-IV based on standards set by the World Health Organization. The TCGA LGG data consist of grades II and III, and the GBM data are considered grade IV. The processing of raw reads alignment, variant QC, ethnicity, and relatedness inference were the same as those used for the CNS and ALL cohorts. After removing related samples, 325 GBM samples and 483 LGG samples remained. We used CoCoRV to perform summary count–based association tests separately for GBM and LGG samples. We selected the non-cancer summary counts from gnomAD as controls and used the CMH ethnicity-stratified test. The threshold of AF and the AF_popmax (the maximal AF among ethnicities from the gnomAD annotation) were set to 1*e*–3. The same criteria were used to define potential pathogenic variants, as in the pediatric CNS and ALL cohorts. We observed no inflation in the QQ plot (Fig. S9). Table 4 shows the top five genes from the results of the GBM cohort. We further tested these five genes in the TCGA LGG cohort and found that two genes, *TP53* and *ABCB8*, were significant (P <0.05) (Table 4). As an independent statistical validation, we used 85 pediatric high-grade glioma WES samples from the PCGP project^15^ as cases and independent 8,525 controls, including additional non-cancer in-house samples and samples from the ADSP and the 1,000 Genomes Project. We used the full genotype–based analysis in this validation for the best control of confounding effects. The top 20 PCs were included as covariates, and the VT method implemented in EPACTS^16^ was used for the rare variant association test. The p-values of *TP53* and *ABCB8* were significant (p=0.01 and 0.005, respectively) (Table 4). *TP53* is a known tumor-suppressor gene and a predisposition gene for many cancers, including adult and pediatric brain tumors^14,17^. Huang *et al*.^5^ performed a germline-based association analysis of TCGA data that was focused on 152 candidate genes; however, *TP53* was not listed as significant for GBM. This false negative is most likely due to their analysis design: they used one cancer versus all other cancers in TCGA, and *TP53* is enriched in multiple cancers so the enrichment in GBM is not detected. *ABCB8* is a novel candidate gene associated with brain tumors that encodes an ATP-binding subunit of the mitochondrial potassium channel. Potassium channels can promote cell invasion and brain tumor metastasis^18^. They also have an important role in brain tumor biology^19^. Per the GTEx portal, the cerebellum has the highest median expression of *ABCB8* (Fig. S8). Further experiments are crucial to confirm whether deleterious *ABCB8* mutations increase the risk to adult brain tumors.

**Table 4.** The top candidate genes from the TCGA GBM cohort and statistical evidence from the TCGA LGG cohort and the PCGP HGG cohort

| TCGA GBM |  |  | TCGA LGG | PCGP HGG |
| --- | --- | --- | --- | --- |
| chr | Genes | P-value | P-value | P-value |
| 1 | *TTC39A* | **7.30e-06** | 1 | - |
| 17 | *TP53* | **5.60e-05** | **0.02** | **0.01** |
| 12 | *ELK3* | **3.80e-04** | 1 | - |
| 7 | *ABCB8* | **4.62e-04** | **0.01** | **0.005** |
| 1 | *CELA2B* | **1.13e-03** | 0.14 | - |

Abbreviations: chr, chromosome; FDR, false discovery rate; GBM, glioblastoma multiforme; HGG, high-grade glioma; LGG, low-grade glioma; PCGP, Pediatric Cancer Genome Project; TCGA, The Cancer Genome Atlas

### Analysis of an amyotrophic lateral sclerosis study using only case summary counts

We used publicly available case summary counts and gnomAD summary counts for rare variant analysis. Specifically, we downloaded the summary counts of an ALS study^20^ from the ALSDB website (http://alsdb.org/downloads). Because we cannot classify the ethnicities of each sample based only on the summary counts, we relied on the reported summary counts of Caucasian ancestry, which included 3,093 cases and 8,186 controls. The data were converted to the VCF format and annotated using ANNOVAR. We used the downloaded coverage bed file to extract high-coverage sites (at least 90% of samples with coverage ≥10) and intersected those with gnomAD high-coverage sites. In addition, we used the UCSC CRG Align 36 track to ensure that the mapping uniqueness was 1. The AF threshold was set to 5e–4. We included variants annotated as stopgain, frameshift_insertion, frameshift_deletion, splicing, and nonsynonymous with REVEL score ≥0.65. We applied our method to the case summary counts and summary counts from the gnomAD nfe population. There was a mild inflation in the QQ plot (Fig. 7a). Known genes *SOD1, NEK1*, and *TBK1* were ranked in the top. Two false positives were confirmed by manual variant QC check. For one false-positive gene, there were eight indels at the same position, chr12:53207583, and only homozygous genotypes were observed. The two false-positive genes had variants in both study-specific cases and controls but none in gnomAD. This observation was most likely due to processing platform–related batch effects. Because study-specific control summary counts were also available, we also performed the summary count–based case-control rare variant analysis using case-control data only from this study. Yet, we still identified no obvious inflation in the QQ plot (Fig. 7b). For both cancer-predisposition genes *NEK1* and *TBK1*, using gnomAD showed a much-improved significance level, compared with that using the study-specific control summary counts. This demonstrates the advantage of using public summary counts of large sample size in prioritizing disease-associated genes. Although minimal variant QC manipulation can be done with only summary counts available from cases and controls in this data set, CoCoRV successfully prioritized the disease-predisposition genes.

**Fig. 7.**
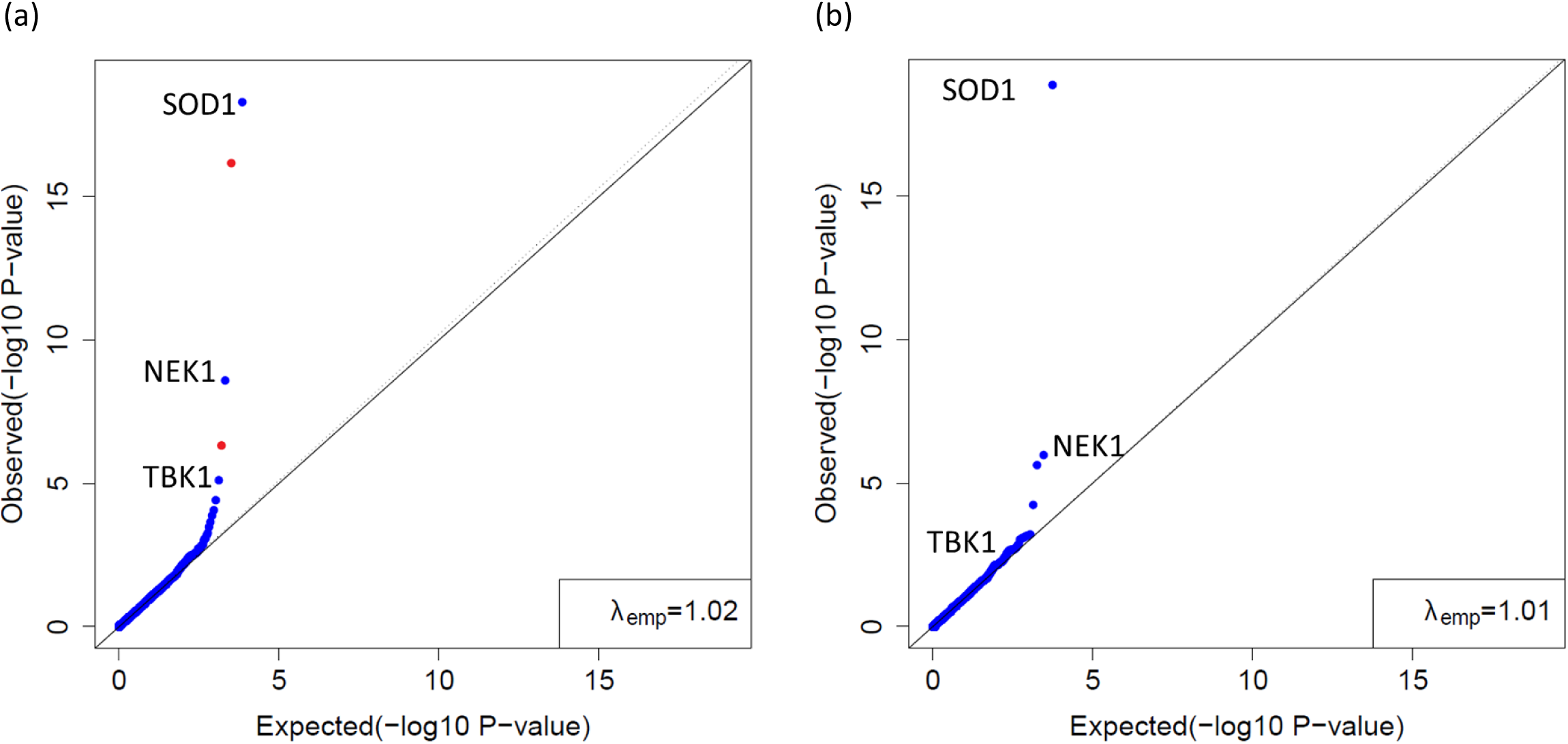
Rare variant analysis of the Caucasian samples from the ALSDB data using summary counts for both cases and controls. a. CoCoRV analysis using control summary counts from the gnomAD non-Finnish Europeans. The red dots are false positives after manual check on the variants contributing to the test statistics: the first false positive is due to a large cluster of indels in the study-specific case and control samples; the second is due to a variant with strong Hardy–Weinberg disequilibrium in both study specific cases and controls. b. CoCoRV analysis using control summary counts from the study specific 8,186 Caucasian control samples.

## DISCUSSION

In this study, we developed and tested a new framework, CoCoRV, that prioritizes disease-predisposition genes by using public summary counts as controls for rare variant burden tests, making it possible to discover novel genes without sequencing additional healthy control samples. Our framework provides consistent variant QC and filtering, ethnicity-stratified gene-based burden test, accurate inflation factor estimation, powerful FDR control, and a novel approach to detect high-LD variants using gnomAD summary counts, which is essential for removing false positives. The inflation factor estimation and FDR control method can also be used in other applications if the tests are performed using FET or CMH. Besides its application in the rare variant association test, our proposed LD test can detect high-LD MNVs, and is not limited by sequencing read length compared with traditional read-based MNV detection. Identifying rare variants in strong LD is also helpful in other analyses, e.g., distinguishing compound heterozygous variants from that on the same haplotype or applying special treatment to MNVs^13^. We also evaluated the concordance between using summary counts and full genotypes. In general, the concordance was good, especially for top-ranked genes with a low p-value threshold.

By applying CoCoRV framework to pediatric and adult cancers, we identified known cancer-predisposition genes and prioritized novel risk genes, one of which was statistically validated. We caution that CoCoRV should be used as a prioritization tool and not a statistical validation tool, because using summary counts might not control some hidden confounding factors. Once interesting genes are identified, either a strict full genotype–based association test or lab-based functional studies are needed to validate the findings.

One potential limitation of using summary counts is that adjusting for covariates is not straightforward. Besides potential batch effects introduced by the sequencing platform, one (and often the only) confounding factor in genetic studies is population structure^21^. We propose to use a CMH-based, ethnicity-stratified analysis to mitigate this problem. Whether the population stratification causes an obvious systematic inflation depends on three factors: 1) population composition in the cases and controls, 2) AF threshold, and 3) sample size. In practice, we cannot guarantee that the populations of cases and controls will match perfectly, but with an adequately low AF threshold (e.g., 5e–4) and a focus on pathogenic variants, the influence of the population structure can be substantially reduced, possibly to the point of no systematic inflation. Improving the adjustment for population structure using summary counts might be worth future investigation.

Testing LD is a well-studied problem in statistical genetics when haplotypes or genotypes can be observed^22,23^. We extended the LD test when only a set of summary counts of two variants was available, such as in gnomAD. We assumed the Hardy-Weinberg equilibrium (HWE) in our test, which simplified the calculation and performed well. This might be due to that only strong LD can be well detected when using a limited set of summary counts, and the influence of the Hardy-Weinberg disequilibrium (HWD) for most rare variants is negligible or inadequate. Further investigation to extend the proposed method to account for the HWD might be useful, for example using a composite measure of LD^24^. Our proposed LD-detection method in CoCoRV is similar to a method proposed to estimate haplotype frequencies and LD measures in pooled DNA data^25^. The major differences are as follows: First, CoCoRV allows for different numbers of samples in a pool, e.g., the subset of samples within each ethnicity in gnomAD, rather than a fixed number in the design of the pooled experiment. Second, instead of using an expectation-maximization algorithm and treating the haplotype frequencies as missing data, our framework uses a direct gradient-based maximization of the likelihood, which exploits many well-developed gradient-based optimization methods^26^; therefore, it converges faster. In addition, we use a one-sided (rather than a two-sided) test to detect strong positive LD. Finally, we use the odds ratio (rather than correlation coefficient) to characterize LD strength for rare variants. In this study, we considered only the LD between two biallelic variants; in the previous study^25^, multiple loci and multiple alleles were allowed, which might be an interesting topic to explore in the future.

One main difference between using the separately called summary counts and jointly called full genotypes was the variant QC step. For example, for the separately called summary counts–based analysis, VQSR was applied separately to cases and controls; however, for jointly called full genotype–based analysis, it was applied to jointly called cases and controls. We found that the results of VQSR for a specific variant can differ between cases and controls if run separately. This is partly expected because VQSR uses a machine learning approach to model the quality of variants within a jointly called genotype matrix. Different studies result in different training samples and different trained models. Including public sequencing data, such as the 1,000 Genomes Project’s WES data, with the case sequencing data will most likely improve the robustness of the variant QC and the concordance when the summary counts–based analysis is performed later.

The lack of obvious inflation in the QQ plot does not guarantee that the top hits are true positives; it indicates only that there is no large systematic inflation in the association test. However, sporadic individual false positives could be among the top hits. When using publicly available summary counts as controls, the cases and controls are processed from different pipelines using different parameters or software versions. Therefore, this approach is more prone to false positives than is the joint analysis of case and controls using full genotypes generated from the same pipeline and QC steps. Also, a more stringent check on the sequence region of the called variants would be helpful, such as alignment uniqueness, duplication segments, or repeats from the UCSC genome browser resources. Warning QC flags from gnomAD are also useful.

The indel variants required more manual inspection due to challenges in the accurate alignment and QC. Another method that alleviates the false-positive problem is to process cases with a relatively smaller, publicly available sequencing data set, such as the WES data from the 1,000 Genomes Project, which is also advocated by GATK. Although this approach requires increased processing time and storage space, it helps separate the true enrichment of rare alleles in cases from the false enrichment due to pipeline differences.

Compared with the standard full genotype–based analysis using the continuous PCs to account for population structure, the coarse assignment of discrete ethnicity groups cannot account for finer population structure within each ethnic group. Accurate inflation factor estimation and the QQ plot is thus critical for determining whether there is systematic inflation due to population structure. For ethnicity stratification, we used the 1,000 Genomes Project’s samples to predict the ethnicity of cases. This procedure was similar to the one used by gnomAD, though the final trained classifier was not the same due to different training set. Hence, some discrepancies may occur between our assignment and those of the gnomAD classifier. We anticipate that the differences would be small, considering that the 1,000 Genomes Project’s samples are also part of the gnomAD data set, and a very high probability threshold (0.9) was used to classify the samples in both our and gnomAD’s ethnicity classification. Moreover, individual small discrepancies in the assignment are unlikely to substantially influence the final rare variant–based association test. When interpreting the association results, it is always more convincing if the association signals contributing to the final significance are found in more than one ethnicity.

In this study, both our processed data and gnomAD summary counts used GATK for variant calling and joint genotype calling, though there were differences, e.g., different versions, detailed implementations, and variant QC. By retaining high-quality variants and maintaining a consistent filtering strategy, we showed that batch effects in the processing pipeline can be well controlled. However, we have not explored the potential batch effects if two completely different variant-calling algorithms were used (e.g., if one uses GATK and the other uses FreeBayes^27^). We anticipate these might introduce some inconsistencies and require further investigation. Recent standardization of some key processing steps in genome-sequencing analysis pipelines^28^ shows promising results in minimizing these batch effects.

## METHODS

### Data sources

The case cohort in our main analyses included pediatric cancer samples from the St. Jude PCGP (HGG), St. Jude LIFE study [central nervous system tumors (CNS) and acute lymphocytic leukemia (ALL)]^29^. We constructed an in-house control cohort of samples using the WES data from the Alzheimer’s Disease Sequencing Project (ADSP)^30^ and the 1,000 Genomes Project^31^, which was later used to compare the concordance between using separately called control summary counts and jointly called case-control full genotypes. The data sets included multiple ethnicities (Fig. S10), with European ancestry being the majority (52%-81%). Inclusion of these individuals in our study was reviewed and approved by Institutional Review Board at St. Jude Children’s Research Hospital. We also tested our framework in two other independent data sets, The Cancer Genome Atlas (TCGA) brain tumor cohort and an amyotrophic lateral sclerosis (ALS) study, to illustrate its power.

### Consistent quality control and filtering of variants

When cases and controls are called separately, coverage summary information (e.g., the percentage of samples with coverage ⩾10) ensures that regions of interest are well covered. Our tool incrementally summarizes coverage information and computes the percentage of samples with coverage no less than specified thresholds, such as {1, 5, 10, 15, 20, 25, 30, 50, 100}, for each nucleotide position. Our tool can scale well for tens of thousands of samples and is easy for parallel running (supplemental text). Following the coverage filtering in Guo *et al*.^4^, we kept the variants that have ≥10 coverage in at least 90% of the samples for both cases and controls.

Inconsistencies can happen when QC and filtering are applied separately, e.g., when a variant is filtered-out from controls but not from cases or *vice versa*. To address this problem, we employed the following strategy: keep only high-quality variants from each cohort’s QC process and perform consistent QC and filtering. For the former, all variants must pass the cohorts’ QC filter. We required that the missingness within cases and controls be ≤0.1. In addition, we used the gnomAD data to generate a blacklist of variants (supplemental text). For consistent QC filtering, we excluded all variants that failed QC steps in either cohort. This consistent filtering step is absent in other tools such as TRAPD^4^.

Another critical filtering step is joint allele frequency (AF)-based filtering. The joint AF of a variant is estimated by pooling the counts of cases and controls together, i.e., *AF =* (*AC*_*case*_+*AC*_*control*_)/(*AN*_*case*_+*AN*_*control*_), where *AC*_*case*_ and *AC*_*control*_ are the alternate allele counts of cases and controls, respectively, and *AN*_*case*_ and *AN*_*control*_ are the total allele counts (including both reference alleles and alternate alleles) of cases and controls, respectively. Using joint AF for filtering avoids inconsistencies when separately filtering variants based on AF within cases and AF within controls.

### Estimation of sample counts in three models using summary genotype counts

We defined three models to count the samples for the burden test (Fig. 1). Let *AC*_*i*_ be the alternate allele count of each variant *i* in a gene for a sample. A sample belongs to one or more of the following models if the sample satisfies the corresponding conditions:

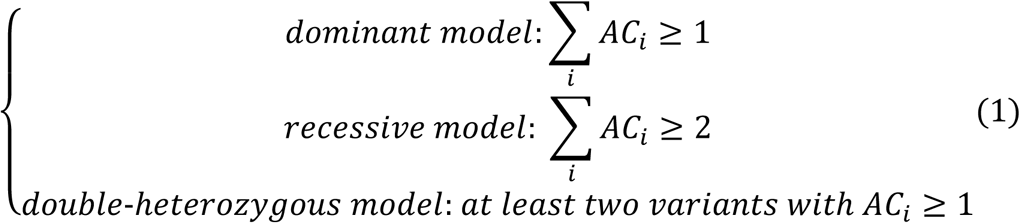

Here the recessive model can be either homozygous with alternate alleles at the same position, or at least one alternate allele at ≥2 positions, i.e., double-heterozygous. Therefore, the double-heterozygous model can be viewed as a special case of the recessive model. As noted in Guo *et al*.^4^, because the haplotype information is not directly observed, the double-heterozygous model could mean two variants on the same haplotype, thus *not exactly* a compound-heterozygous model. We often have the full genotypes for cases, so we can determine the count of each model directly from the genotype matrix. However, for controls, we need to estimate the number of samples qualified for each model based on the summary counts of each variant. Specifically, for a gene with *m* variants, let *p*_*iG*_ be the genotype frequency of variant *i* with genotype *G*, where *i=* 1, ⋯*m*, and *G =* 0, 1, 2. Public summary counts, such as gnomAD, usually provide this information; however, if only AF is available, the genotype frequencies must be estimated. One convenient, though not optimal, approach is to assume the Hardy Weinberg equilibrium (HWE). We estimate the probability of a sample satisfying the defined models in (1) as follows. Denote the probability of the dominant model by *p*_*DOM*_, the recessive model by *p*_*REC*_, and the double-heterozygous model by *p*_2*HET*_. If only one variant exists, we have *p*_*DOM*_ = 1 − *p*_10_, *p*_*REC*_ = *p*_12_, and *p*_2*HET*_ *=* 0. If there are at least two variants, they can be estimated as shown below, assuming independence between the rare variants:

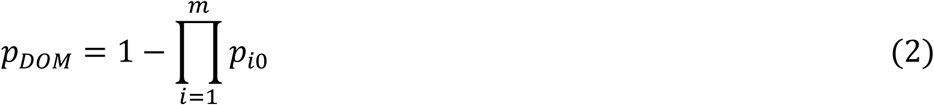

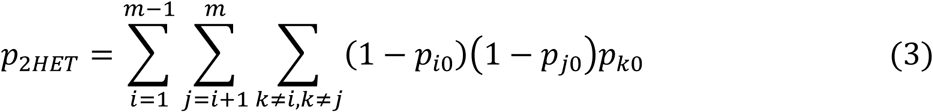

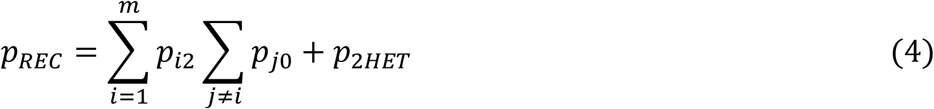

Equation (2) calculates the probability of being dominant for at least one locus. For (3), we approximated the probability by considering all pairs of variants with *AC* ≥ 1 out of *m*loci, because the probability of having two variants with *AC* ≥ 1 is much higher than that of having three or more variants with *AC* ≥ 1. Similarly, in (4), we considered a single variant with homozygous alternate alleles and two variants with *AC* ≥ 1. To calculate the estimated counts, we multiplied the frequencies of each model by the total number of controls. Here we assumed all rare variants were independent; later we made some modifications when we detected variants in LD.

### Burden test with samples stratified by ethnicity groups

In addition to the pooled analysis of all ethnicities, we provide ethnicity-stratified analysis. The latter requires predicting the ethnicity of each case. We performed principal component (PC) analysis for all samples, including samples from the 1,000 Genomes Project (see the supplemental text). Following the practice in gnomAD^1^, we trained a random forest classifier based on the top 10 PCs and used the six major ethnicity categories as training labels: non-Finnish European (nfe), African American (afr), Admixed American (amr), East Asian (eas), South Asian (sas), and Finnish from the 1,000 Genomes Project. Then we applied the classifier to the rest of the samples and used probability 0.9 as a cutoff to assign samples to the six ethnicities or to “other” when all predicted probabilities were less than 0.9. We then stratified samples based on the six ethnicities and calculated/estimated the counts for each model within each stratum as described above; samples labeled “other” were not used. We performed the CMH-exact test^32^ to aggregate evidence of each stratified 2×2 contingency table. When using gnomAD summary counts as controls, we used the stratified summary counts of each ethnicity directly.

### Inflation factor estimation for discreate count based test

For the p-values of burden tests calculated by the FET or CMH, the null distribution was not the uniform distribution. Therefore, the expected ordered null p-values could not be derived from the uniform distribution. The null distribution of the ordered p-values of a set of tested genes using either FET or CMH depends on the number of rare alleles in each gene. Instead of assuming the incorrect uniform distribution, we empirically sampled from the correct null distributions for inflation factor estimation. We assumed that the genes are independent of each other, which is reasonable for rare variants with very low AFs. Instead of permuting the phenotypes, we worked on the 2×2 contingency table directly (Fig. 2a). For each gene, let *m* be the number of cases, *n* be the number of controls, and *k* be the number of samples with rare alleles among all cases and controls. These values can be obtained from the observed 2×2 contingency tables and are held fixed for each gene. Then we sampled the number of case samples with rare alleles (denoted by *x*) from a hypergeometric distribution with parameters *m, n, k* and formed a new 2×2 contingency table under the null hypothesis of no association. We sampled the contingency table for each gene by using gene-specific parameters *m, n, k* to generate 2×2 contingency tables for all genes. Then we obtained the null p-values using FET and sorted them across all genes. This process was repeated *N* times, and the final expected sorted p-values (order statistics) were the average of *N* p-values at each ordered rank. To estimate the inflation factor, we took the lower 95% quantile of points in the QQ plot and regressed the sorted log_10_-scaled observed p-values to the log_10_-scaled expected sorted p-values. The slope of the regression was used as the inflation factor, which is denoted by *λ*_*emp*_. Similarly, we extended the inflation factor estimation to the CMH-exact test. To simulate the *empirical* p-values under the null hypothesis for each stratified contingency table, we sampled a new 2×2 contingency table as described above and then obtained the null p-values by using the CMH-exact test. The inflation factor could then be calculated similarly as in FET.

Guo *et al*.^4^ also noted an excess of p-values that equaled 1 when the null distribution was assumed to follow a uniform distribution. Thus, they proposed modifications to estimate the inflation factor. Two versions of this estimation are included in Guo *et al*.^4^. One estimated the slope by using values between quantile 0.5 and 0.95, denoted by *λ*_0.5−0.95_ in our study, and the other estimated the slope from two points: the first is the point with the observed p-value of 1 with the highest rank, and the second is the point at the 0.95 quantile of the p-values not equal to 1, denoted by *λ*_*2points*_ in our study. For both *λ*_0.5−0.95_ and *λ*_2*points*_, the expected ordered p-value of rank *r* in an increasing order from *n* p-values was *r*/(*n*+1) assuming a uniform distribution *U*(0, 1). Because these modifications were still based on a uniform distribution, their solutions were biased.

To illustrate the performance of different methods on inflation factor estimation, we used simulated p-values under the null hypothesis to evaluate the bias. Specifically, for *N* replicates of sampled null p-values, we applied all three methods to estimate the inflation factor. For the empirical null p-value–based method, we saved on computation cost for each replicate of null p-values by using the rest *N* − 1 replicate of null p-values to estimate the expected sorted p-values.

### Resampling based FDR control

Resampling based FDR control method was originally developed to address the correlation between multiple tests^33^. We adopt it here for FDR control of discrete count based tests. We simulated the p-values under the null hypothesis the same as in the inflation factor estimation (Fig. 2a). Then we adopted the resampling based FDR control method to estimate the adjusted p-values. The two resampling methods used a mean point estimate (RBH_P) or an upper limit estimate (RBH_UL) for the number of true positives detected, respectively. Given the simulated p-values under the null hypothesis, the adjusted p-value estimation is similar as that in the R package FDR-AME^34^. Let *R*^*∗*^(*p*) be the number of genes with p-values no greater than *p* in the simulated p-values under the null hypothesis, *R*(*p*) the number of genes with p-values no greater than *p* in the observed p-values, *M*^*∗*^(*p*) the mean of 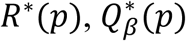 the 1 − *β* quantile of *R*^*∗*^(*p*). Then for RBH_P, the estimated true positives detected at threshold *p* was *s*(*p*) *= R*(*p*) − *M*^*∗*^(*p*); for RBH_UL, 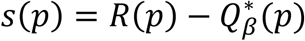. Then the adjusted p-values were calculated as 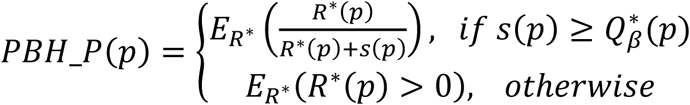 and 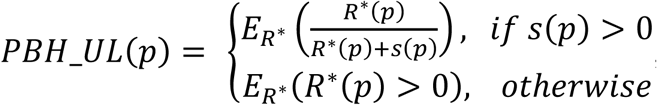, where *E*_*R*_*∗* means taking the expectation over all replicates of p-values under the null hypothesis.

### Detection of high-LD rare variants via gnomAD summary counts

One interesting feature of CoCoRV is the proposed LD test using only gnomAD summary counts (Fig. 2). The gnomAD data set has several subset-based summary counts, including “controls” and “non_cancer.” Because control samples are a subset of non_cancer samples, we can partition them into three independent sets of summary counts: cancer, healthy controls within the non_cancer set, and other diseases within the non_cancer set, which are labelled “non_cancer_non_controls.” After further stratifying by sex, we can generate six independent summary counts *per* ethnicity (Fig. 2). We assume that the allele frequencies and LD among variants are the same among these six independent data sets per ethnicity, which is likely reasonable because most variants should not be associated with sex or cancer. Given the *ACs*(alternate allele counts in a cohort) of two variants and the total haplotypes in these data, we can test the hypothesis that two variants are in positive LD, i.e., the variants are more likely to lie on the same haplotype than random chance under the assumption of independence. Specifically, let the data observed be {*x*_*i*_, *y*_*i*_, *n*_*i*_, *i=* 1 ⋯*I*}, where *x*_*i*_is the *AC* of the first variant, *y*_*i*_ is *AC* of the second variant, *n*_*i*_ is the total number of haplotypes, *I* is the number of independent sets of summary counts (*e*.*g*., six from gnomAD) (Fig. 2). Denote the four haplotypes of two variants by *h*_11_, *h*_10_, *h*_01_, *h*_00_ and their corresponding probabilities by *p*_11_, *p*_10_, *p*_01_, *p*_00_, where 1 indicates the alternate allele and 0 indicates the reference allele. The log likelihood of the observed data is

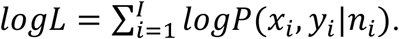

In the following equation, we drop the subscript *i* to simplify the notation. Let *r* be the count of unobserved haplotype *h*_11_, then the counts of the four haplotypes are *r, x* − *r, y* − *r*, and*n*− *x* − *y* +*r*, respectively. By the law of total probability, the likelihood of observed allele counts *x, y* of two variants given *n* total haplotypes is *P*(*x, y*|*n*) *=*∑_*r*_*P*(*x, y, r*|*n*). Assuming the HWE, *P*(*x, y, r*|*n*) can be calculated as the multinomial probability with the four haplotype counts and probabilities. Specifically,

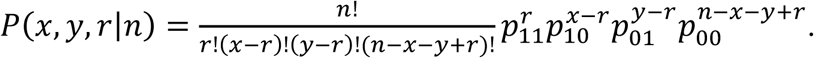

Note that the range of *r*is [max(0, *x* + *y* − *n*), min(*x, y*)]. There are only three free parameters for the four haplotype probabilities because their sum is 1. For a direct test of the null hypothesis, we reparametrize the haplotype probabilities using another three parameters *s= p*_11_ + *p*_10_, *t= p*_11_ +*p*_01_, and 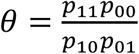, where *s* is the alternate AF of the first variant, *t* is the alternate allele frequency of the second variant, and *θ* is the odds ratio specifying the LD strength and direction between two variants. We used a likelihood ratio test, where the null hypothesis was *θ* ≤ 1 and the alternative hypothesis was *θ* > 1, i.e., the two variants were more likely to be on the same haplotype. The parameters *s* and *t* were treated as nuisance parameters. Specifically, the test statistic is as follows:

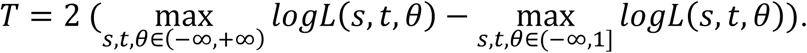

We derived the gradient of the log-likelihood function and used the R package *nloptr*^26^ to maximize the log likelihood under the null hypothesis and full model. The chi-square distribution with 1 degree of freedom was used to calculate the p-values, which appeared to work well, though the asymptotic distribution of the one-sided likelihood ratio test could be better characterized. We used an odds ratio of 1 in the null hypothesis; however, other prespecified odds ratios can be used to test directly whether the LD exceeded a high odds ratio threshold. Due to QC, the *n*_*i*_ of each variant can slightly differ, hence in our implementation, it was calculated as the rounded average of all (*n*_*i*_)s of the qualified variants within a gene. To accelerate the optimization, we implemented the gradient functions in C++ with Rcpp^35^.

The above test can also be used to detect LD when full genotypes are observed. The additive coding of genotypes corresponds to *n* = 2 in the above summary count–based test. The calculation of the likelihood can be accelerated because there are only nine combinations of genotypes between the two variants. Let *f*_*ij*_*= P*(*x = i, y = j*|*n=* 2), then assuming HWE we have

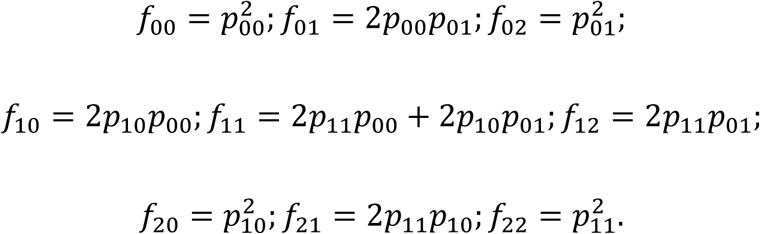

The log likelihood of the data is as follows:

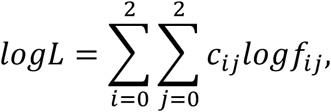

where *c*_*ij*_ is the count of genotype combination *x = i, y = j*. This full genotype–based LD test resembled a recently proposed LD test^23^, but we used a different parametrization. We also implemented this full genotype–based LD test in CoCoRV, in case users have full genotypes, which will result in higher power to detect LDs.

### Accounting for LD between two variants by using gnomAD summary counts

For variants in high LD, we needed to adjust the procedure of counting qualified samples with rare alleles. When estimating the counts of qualified samples in controls for each group of variants in high LD, only the variant with the highest AF was kept; the rest were excluded. For a quick check of high-LD variants, we precomputed the LD test results (p-values and FDRs) from gnomAD and stored those with relatively small p-values (e.g., p <0.1). When estimating sample counts for controls, we regarded variants as high-LD variants if the LD test result has FDR <0.05. To be conservative, for each double-heterozygous sample in the cases, we further required the p-value of the LD test from gnomAD to be less than a threshold (e.g., p <0.05 or 0.1). This practice reduced false positives due to strong LD between variants under the recessive or double-heterozygous models. Note that this processing mainly aimed at variants in high LD, which is the main source of false positives in the association test. Further refinement considering medium or low LD is left for future work.

### Detection of high-LD variants in gnomAD exomes

We used our proposed LD-detection method to scan the gnomAD exome–based summary counts in each gene to detect high-LD variants, which share some similarities with MNV^13^. MNVs are usually detected by direct examination of the sequence reads in aligned bam files. As a validation of the effectiveness of the proposed test, we detected high-LD variants by using the summary counts from the gnomAD exomes and compared them with the reported gnomAD MNVs.

Because we are interested in rare variants annotated with some functions, we focused on variant sets annotated as “stop gain”, “nonsynonymous”, “splicing”, “frameshift_insertion”, or “frameshift_deletion” by ANNOVAR. We restricted variants in the cohort to those with AF ≤0.01, at least 10 alternate allele counts (*AC* ≥10), and missingness ≤0.1. Note that the alternate allele count differed from the alternate read count, which is the number of sequencing reads harboring the alternate allele supporting the genotype calls. We filtered-out variants in the blacklist, as described in the supplemental text. We required a coverage depth of at least 10 for 90% of the samples. For each pair of variants within a gene, we applied the LD test by using the summary counts for each ethnicity. In total, about 10 million tests were run. Variant pairs with FDR <0.05 within each ethnicity were considered significant. For comparison with gnomAD MNVs, we downloaded the coding MNVs detected from gnomAD exomes, where variants’ distances were ≤2 base pairs (BPs). Therefore, we restricted our comparison to high-LD variants with distances ≤2 BPs. We also downloaded the MNV lists based on gnomAD genomes and defined the union of coding MNVs and genome MNVs with distances ≤2 BPs as the full set of MNVs reported in gnomAD with ≤2 BPs.

### Comparison of CoCoRV with TRAPD

Because TRAPD^4^ was developed to use gnomAD summary counts, we compared it and CoCoRV, with both using gnomAD summary counts. The schematic diagram of applying TRAPD is shown in Fig. S11. TRAPD proposed to use a single annotation QD for variant QC. We followed the description in TRAPD^4^ to select the best QD threshold pairs for cases and controls by which TRAPD’s estimated inflation factor was closest to 1. Because TRAPD uses the maximal allele frequency among different ethnicities (AF_popmax) to filter variants, we added AF_popmax to filter variants in CoCoRV. Specifically, we used an AF_popmax threshold of 1*e*–3 in TRAPD for filtering in the final test with the best QD threshold for single-nucleotide variants (SNVs) and indels. For CoCoRV, we used an AF threshold of 1*e*–3 and AF_popmax threshold of 1*e*–3. Because TRAPD uses only *AC* and *AN* to derive qualified sample counts, we included the analysis using only *AC* and *AN* from gnomAD for CoCoRV. In addition, we included the ethnicity-stratified results for CoCoRV. We changed the default one-sided FET to a two-sided FET in TRAPD to match that used in CoCoRV.

## Data Availability

ALS data: http://alsdb.org/downloads

Alzheimer’s Disease Sequencing Project (ADSP): https://www.ncbi.nlm.nih.gov/projects/gap/cgi-bin/study.cgi?study_id=phs000572.v8.p4

1,000 Genomes Project: https://www.internationalgenome.org/

The Cancer Genome Atlas (TCGA): https://www.ncbi.nlm.nih.gov/projects/gap/cgi-bin/study.cgi?study_id=phs000178.v11.p8

St Jude cancer cohort: https://www.stjude.cloud/

## Code Availability

The code for summary counts–based rare variant association test are available at: https://bitbucket.org/Wenan/cocorv/src/master/

## Acknowledgements

This study was supported by the National Cancer Institute grant P30 CA021765 and American Lebanese Syrian Associated Charities (ALSAC). The content is solely the responsibility of the authors and does not necessarily represent the official views of the National Institutes of Health. We acknowledge the permission granted to use data from the Alzheimer’s Disease Sequencing Project (dbGaP Study Accession *phs000572*.*v8*.*p4*) and The Cancer Genome Atlas (dbGaP Study Accession *phs000178*.*v11*.*p8*). We acknowledge Michael Edmonson’s help with the conversion between plain text–based matrix data format and the VCF format and Angela J McArthur’s helpful scientific editing.

## Author Contributions

W.C. proposed the study design, developed methods, implemented methods, processed data and performed data analysis, review the results and drafted the manuscript. S.W. implemented methods, processed data. S.S.T. implemented methods, processed data. D.E. provided guidance on phenotype subtype information and reviewed the results. G.W. proposed the study design, processed data, reviewed the results and drafted the manuscript. All authors read and approved the final manuscript.

---

**Fig. S1.**
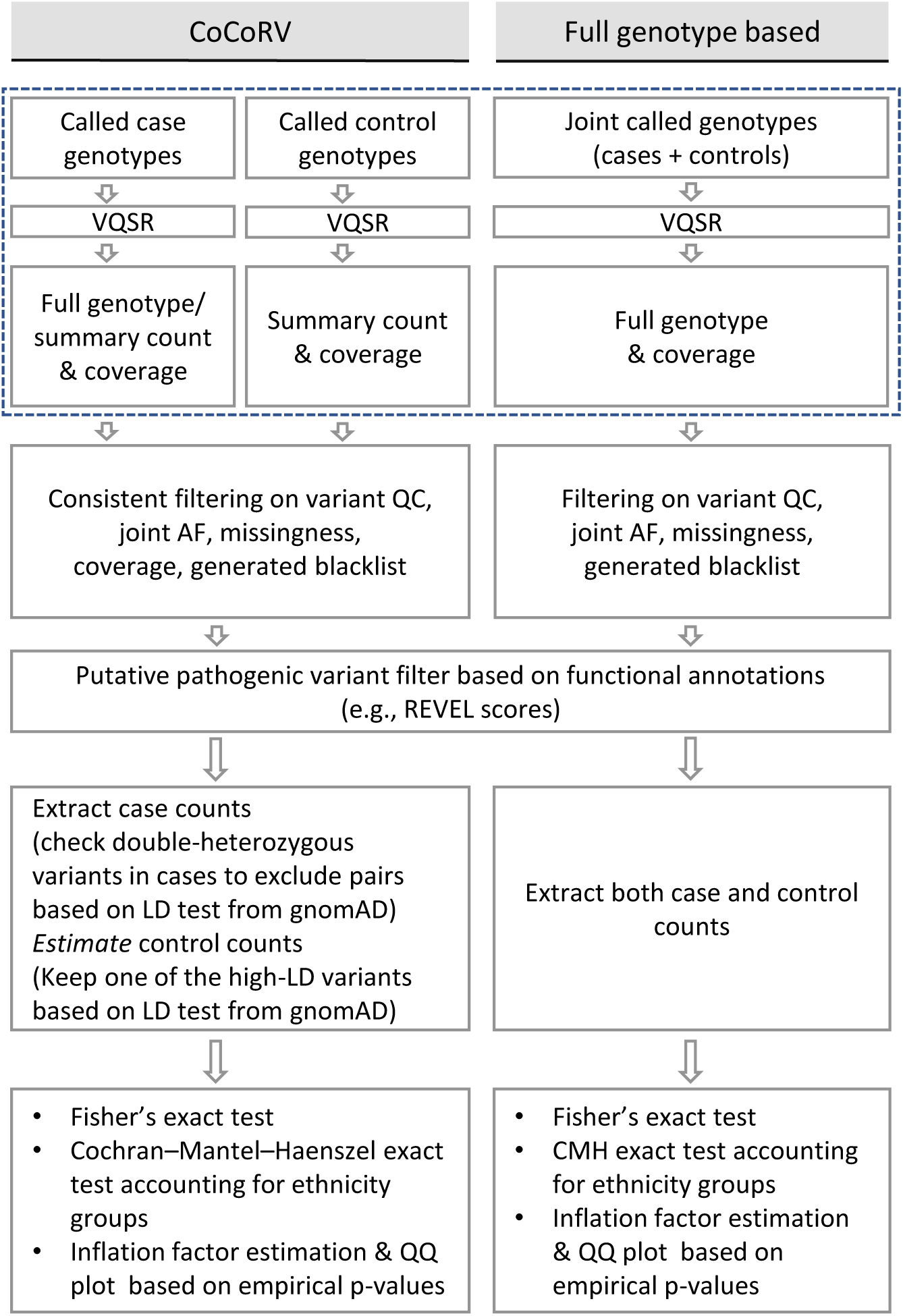
Schematic diagrams comparing the proposed CoCoRV framework using summary counts and jointly called full genotype–based analyses. The left panel shows the CoCoRV framework using summary counts of separately called control cohort. The right panel shows the processing framework when cases and controls are jointly called. The diagrams within the dashed box show how data are generated differently for each analysis framework.

**Fig. S2.**
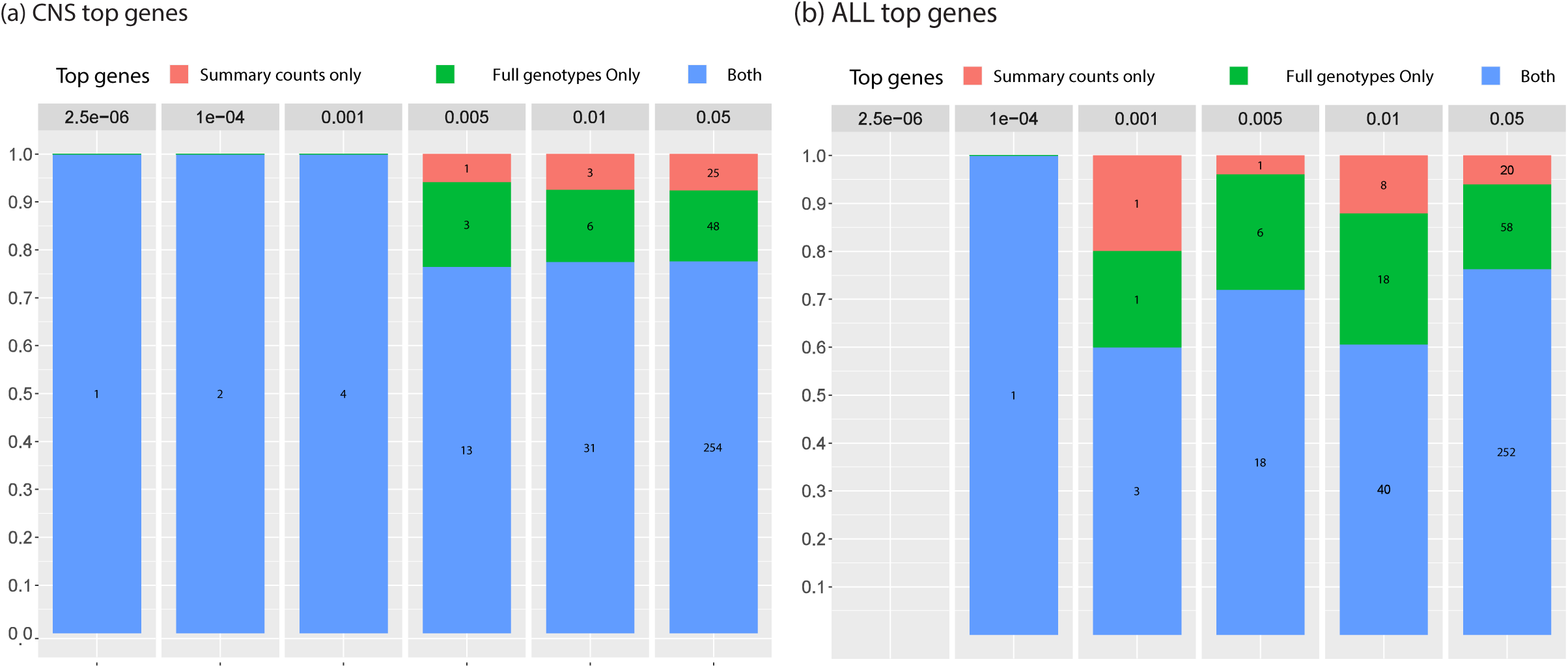
Comparison of top genes between analyses using jointly called full genotypes and separately called summary counts using CoCoRV with the CMH exact test stratifying samples by ethnicities. The bar heights show the percentage and the numbers within each bar show the absolute number of genes. The p-value thresholds used are shown at the top. Results are based on analyses of the CNS (a) or ALL (b) cohort

**Fig. S3.**
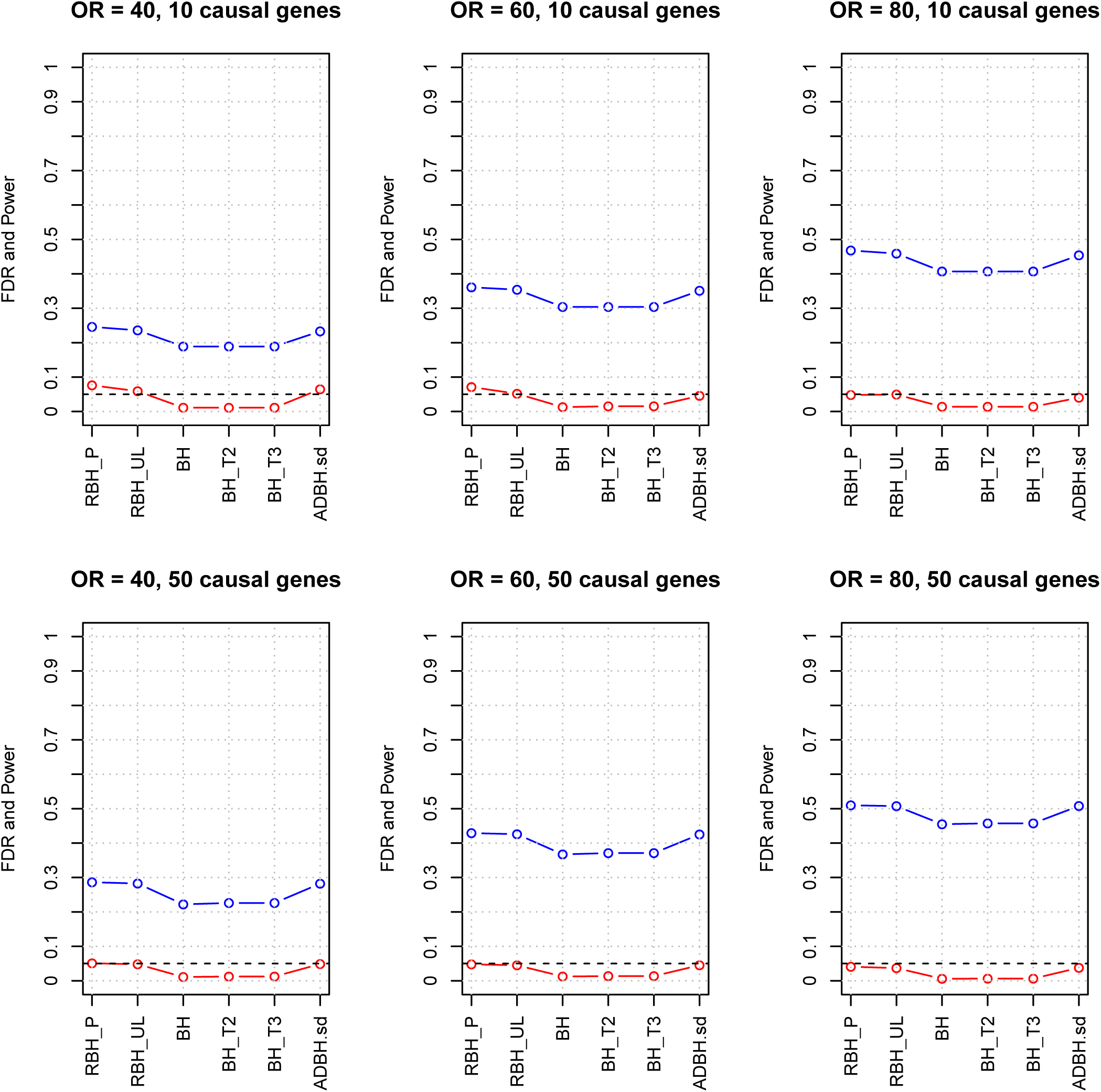
Empirical FDR and power of different methods under different simulation settings. The performance under different combinations of the odds ratio (OR) and the number of causal genes was evaluated. The red color indicates empirical FDRs, the blue color indicates empirical powers. RBH_P: resampling based p-value adjustment using the point estimation; RBH_UL: resampling based p-value adjustment using the upper limit estimation; BH: the Benjamini Hochberg FDR control procedure; BH_T2, BH_T3: methods that remove genes with rare allele counts less than 2 or 3 and then apply the BH procedure; ADBH.sd: A-DBH-SD method from the R package DiscreteFDR.

**Fig. S4.**
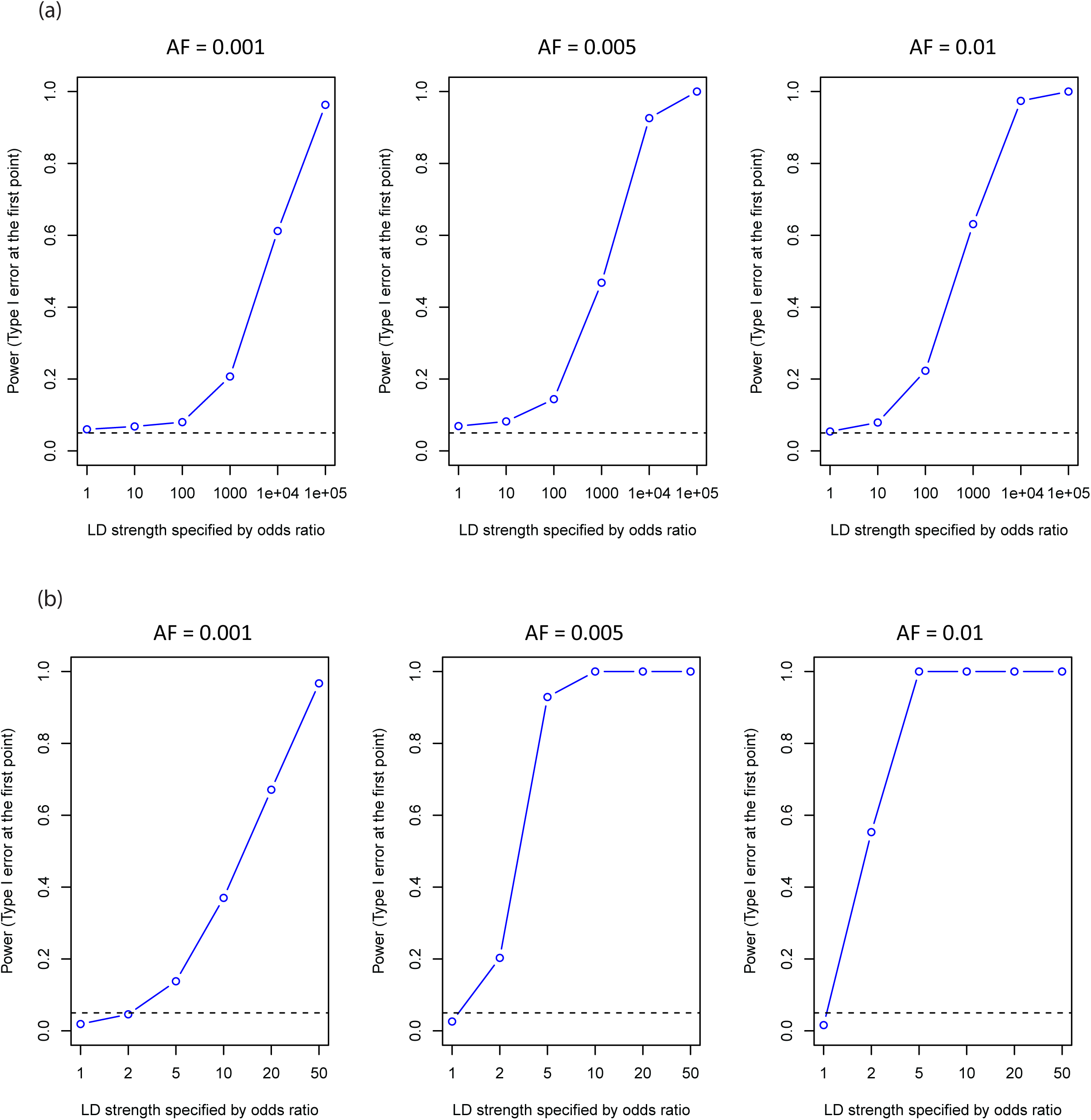
Type I error and power of the proposed LD detection. a. Type I error and power of detecting LDs between rare variants based on six independent groups of summary counts. b. Type I error and power of detecting LDs between rare variants based on full genotypes. Type I error corresponds to the power when the odds ratio is 1. The nominal p-value threshold is 0.05, as indicated by the horizontal dashed line. AF, alternate allele frequency.

**Fig. S5.**
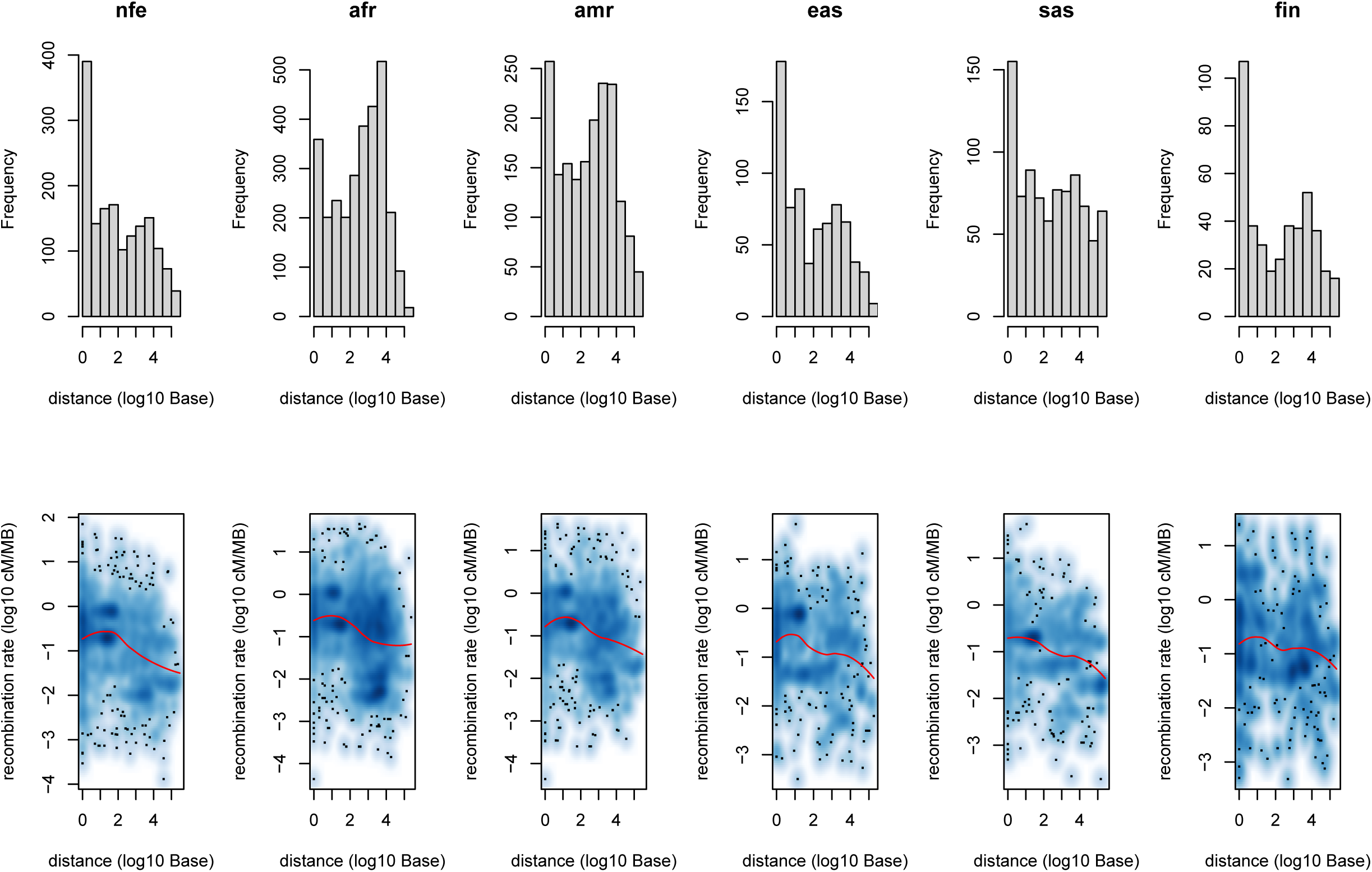
Histograms of the distances of detected high-LD variants in gnomAD and the scatter plot of recombination rates versus the base-pair distances of detected high-LD variants in gnomAD. The distance between two variants is calculated based on the variants’ positions after variant normalization. The upper panel shows the histograms of distances among different ethnicity groups. The lower panel shows the density of scatter plots between the recombination rates and the distances. The recombination rate is based on the middle point between two variants, and it is calculated based on linear interpolation using the genetic map downloaded from the Impute2 website (https://mathgen.stats.ox.ac.Uk/imputeAmpute_v2.html#reference). The red line shows the nonlinear LOWESS fit. Abbreviations: nfe, non-Finnish European; afr, African American; amr, Admixed American; eas, East Asian; sas, South Asian; fin, Finnish; cM, centimorgan; MB, mega base-pairs.

**Fig. S6.**
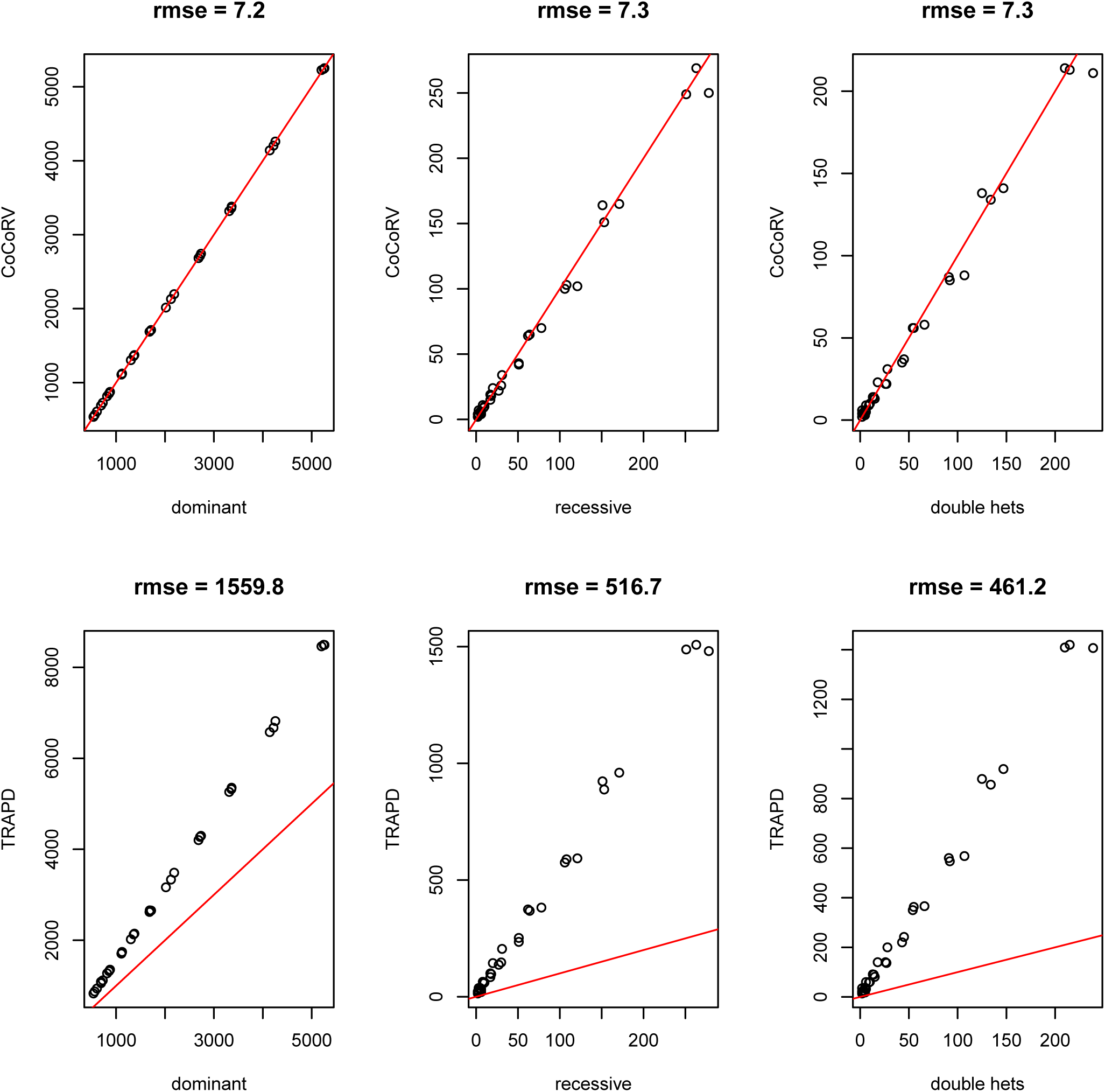
Comparisons of count estimations under different models between CoCoRV and TRAPD. Top panels show the estimates from CoCoRV under the three different models (dominant, recessive, and double-heterozygous), and the bottom panels show those from TRAPD under the three models. The red lines are the y = x lines; rmse, root mean squared error.

**Fig. S7.**
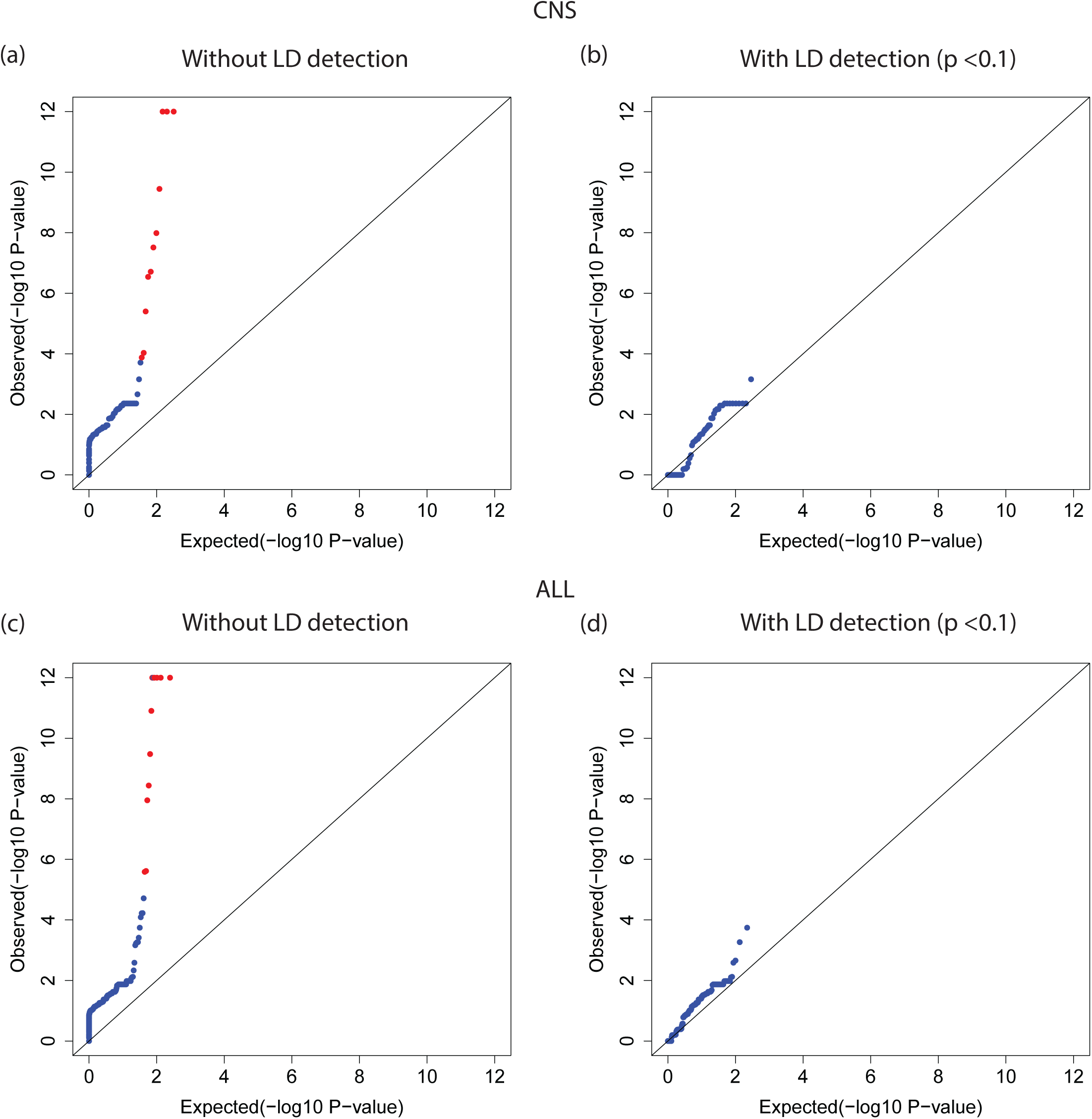
Employing LD detection removes false positives under the double-heterozygous model. a, c. QQ plots of association tests under the double-heterozygous model without employing an LD test in the CNS (a) and ALL (c) cohorts. b, d. QQ plots of association tests after employing LD detection with a p-value threshold of 0.1 in the CNS (b) and ALL (d) cohorts.

**Fig. S8.**
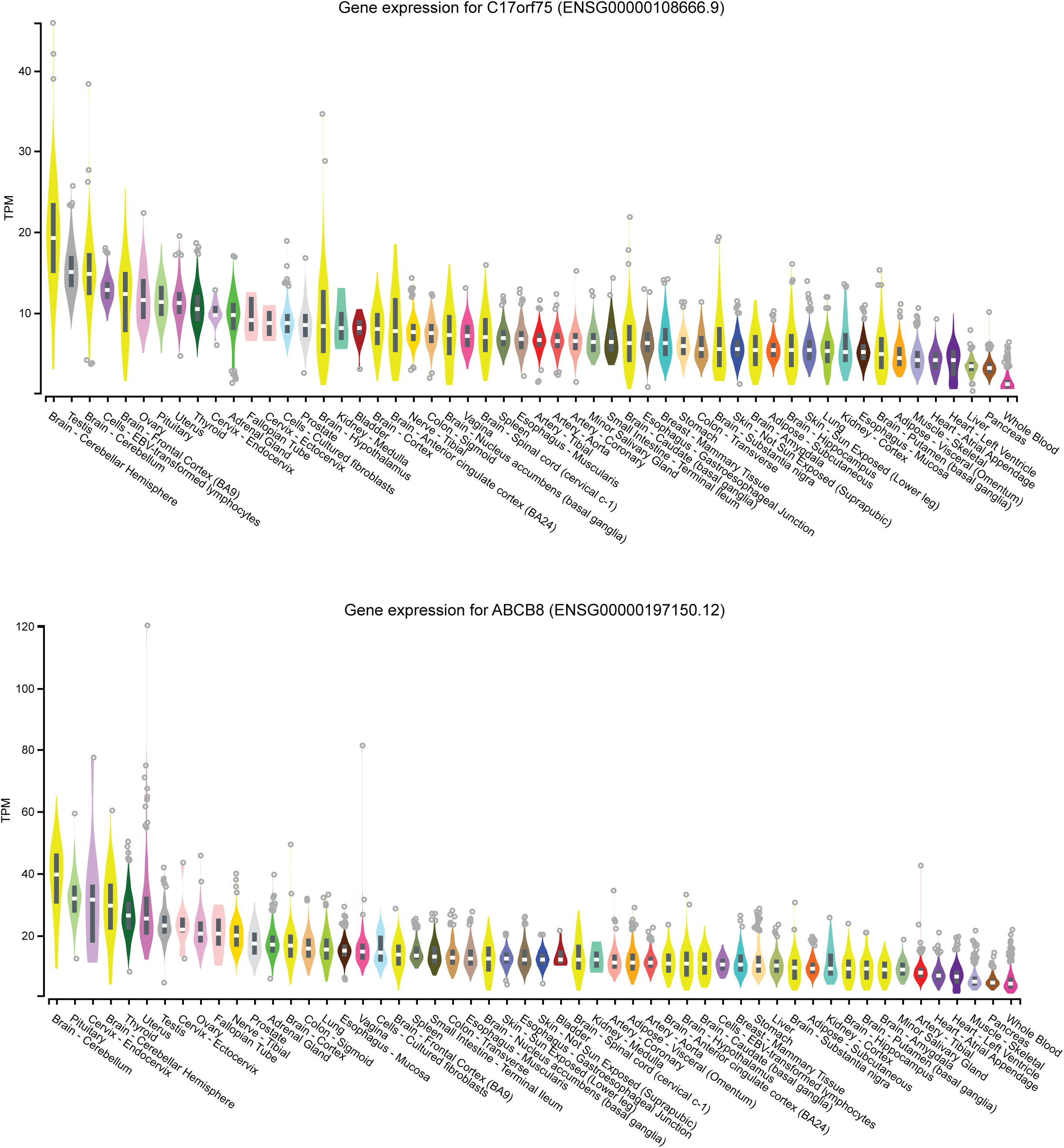
The expression of C17orf75 and ABCB8 genes in different tissues from GTEx. Data obtained from the GTEx portal.

**Fig. S9.**
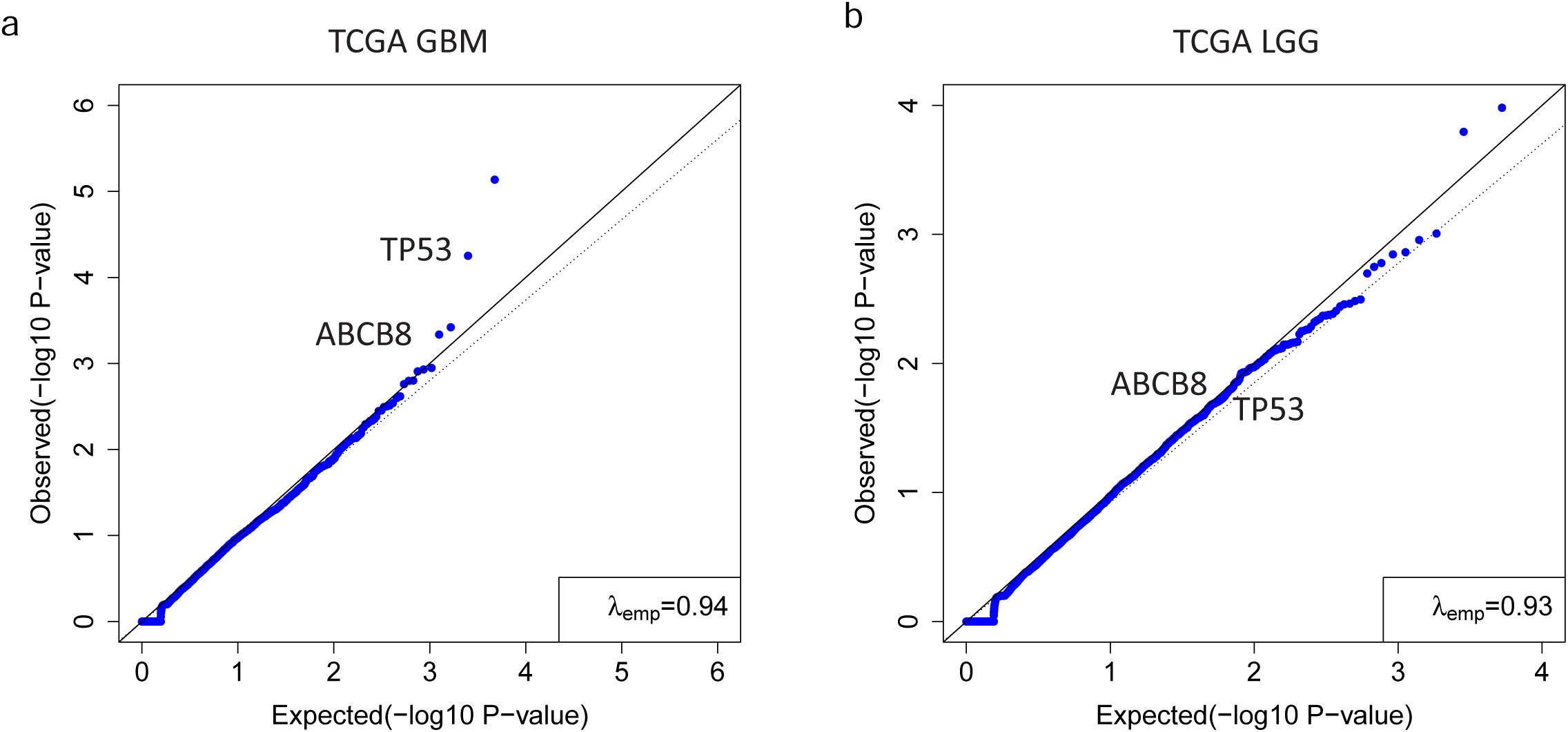
QQ plots of the association tests for the TCGA GBM cohort and TCGA LGG cohort using the CoCoRV framework with gnomAD summary counts as controls. a. QQ plot of the association results of the TCGA GBM cohort. b. QQ plot of association results of the TCGA LGG cohort.

**Fig. S10.**
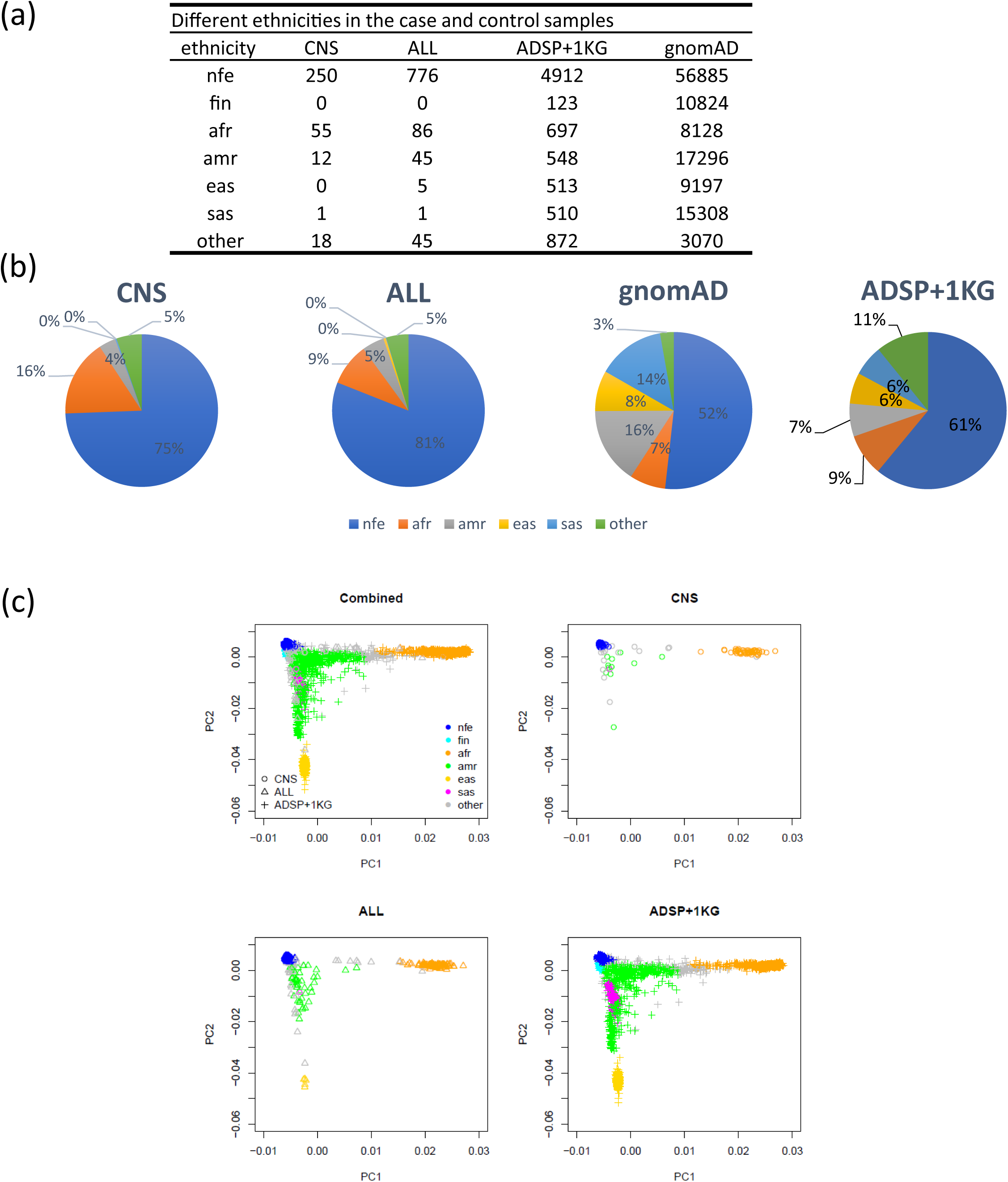
Population structure of the data in our main analyses. Data sets included the CNS and ALL pediatric cancer cohorts and constructed in-house controls with samples from the Alzheimer’s Disease Sequencing Project (ADSP), the 1,000 Genomes Project (1 KG), and the gnomAD. a. The number of samples in each ethnicity, b. Pie charts showing the percentage of each ethnicity in each data set. c. Plots of the top two principal components of the CNS, ALL, and in-house controls. Abbreviations: nfe, non-Finnish European; afr, African American; amr, Admixed American; eas, East Asian; sas, South Asian; fin, Finnish; ALL, acute lymphoblastic leukemia; CNS, central nervous system cancer

**Fig. S11.**
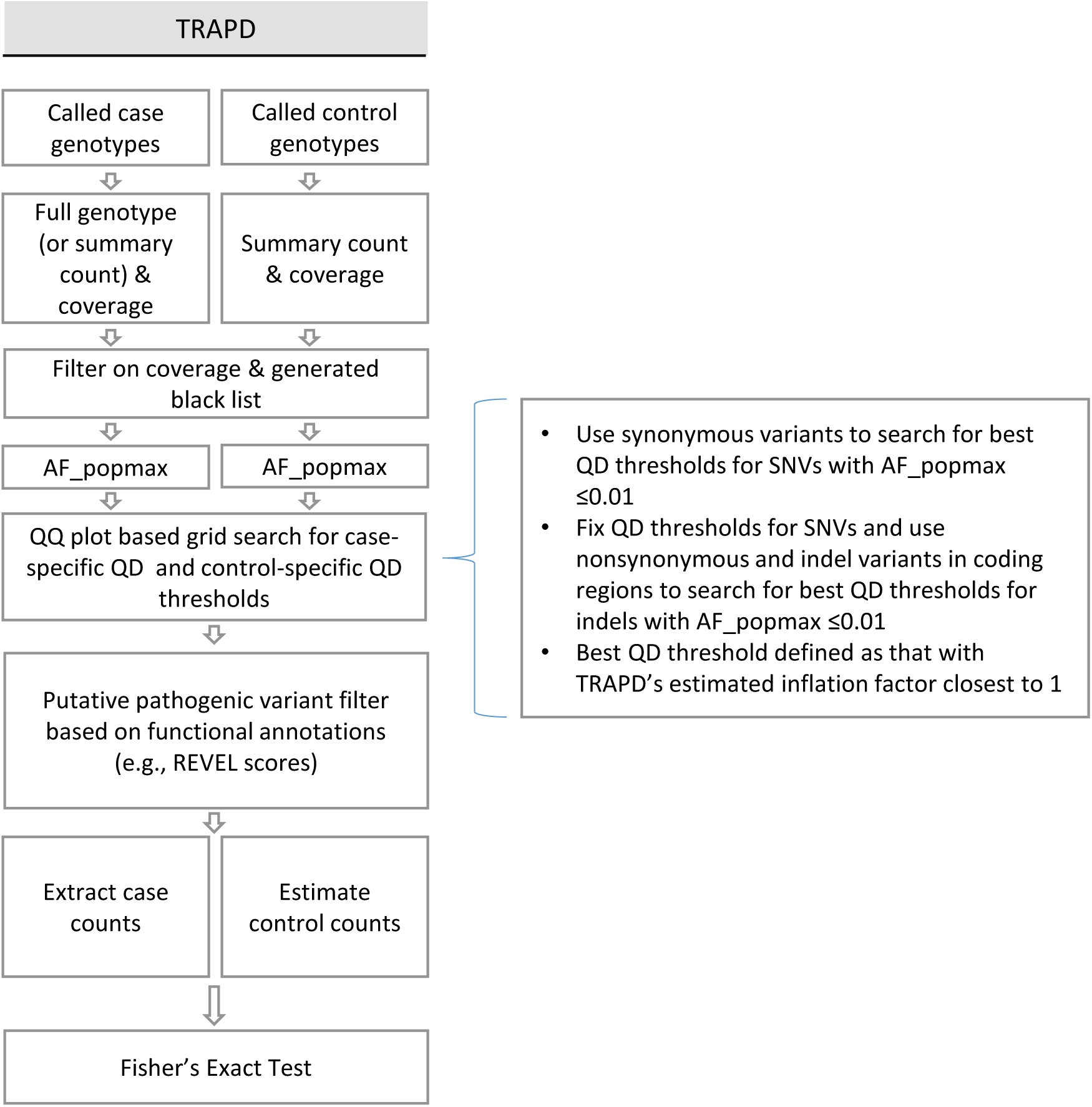
Diagram of TRAPD analysis applied in the study.

