## Supplemental text for "CoCoRV: a rare variant analysis framework using publicly available genotype summary counts to prioritize germline disease-predisposition genes"

### S1 Text. Supplemental Methods

#### Efficient calculation of coverage summary information for subsequent coverage-based filtering

We used samtools[1] to calculate the coverage depth at all qualified positions. We used the aligned bam input, with the minimal base quality ≥10 and the minimal mapping quality ≥20, following the coverage calculation in gnomAD[2]. Each sample was processed independently; therefore, this step can be easily parallelized for each sample. Then we aggregated the coverage data of each sample incrementally to update the summary information, e.g., the total coverage and the total number of samples with coverage depth ≥*x*, where *x* is from the set $\{1, 5, 10, 15, 20, 25, 30, 50, 100\}$. At the end, the summed numbers were divided by the total number of samples to calculate the average coverage and the proportions of samples greater than or equal to each coverage threshold. Instead of simultaneously calculating the depth at each position for all samples, e.g., using GATK *DepthOfCoverage*, this design of calculating the coverage summary information can be easily scaled-up to a large number of samples with constant memory requirement and easy parallelization.

#### Variant blacklist from the gnomAD

The publicly available gnomAD database has a very large sample size and has undergone comprehensive quality control (QC) analysis[2]. For example, it employs a random forest-based and allelic-specific QC to achieve better performance than VQSR. Therefore, we used gnomAD filtering status (gnomAD v2.1) for further QC to remove likely false positives. The following 3 types of variants were excluded: 1) those that failed in both gnomAD whole-exome sequencing (WES) data and whole-genome sequencing (WGS) data; 2) those that failed in one platform and had no variant in the other; and 3) those with an alternate allele frequncy (AF) that differed substantially between WES and WGS. The last was done by testing the null hypothesis that the AF is the same within the WES and WGS in each of the five ethnicities in gnomAD (nfe, afr, amr, eas, fin). The test was done both individually within each ethnicity using a FET if not feasible, a chi-square test was used) and a combined test using a CMH test. Considering the potential differences in population structure between WES and WGS, we required that the p-value from the CMH test be <1e–15. Also, for at least one population, the p-value of the individual test should be <1e–10, and the ratio between AFs from the WES and WGS should be >1.2/1 or <1/1.2, or the absolute AF difference should be ≥0.15.

#### Principal component analysis and relatedness estimation

We used high-quality variants to calculate the principal components (PCs) and estimate the relatedness. Specificly, we used plink v1.9[3,4] to keep high-quality variants with GQ ≥20 and DP ≥10. Then we filtered variants with missingness ≤0.01 and MAF ≥0.001. Samples with missingness >0.25 were also removed. Variants were pruned using plink with the option "--indep-pairwise 1000 50 0.1." Then we used *pcair* and *pcrelate* from the R package GENESIS v2.4.0 [4] to calculate the PCs and estimate the relatedness.

##### Simulation to assess performance of FDR control methods accounting for discrete counts

We simulated 10 or 50 causal genes under the alternative hypothesis and 20,000 genes under the null hypothesis. We set the number of cases to 300 and the number of controls to 50,000. We used the Beta(1, 10,000) distribution to simulate the frequency of pathogenic mutations in one copy of a gene and only kept the rare frequencies, specifically $<5\times10^{-4}$. The total number of samples with pathogenic mutations was then set by the simulated frequency multiplied by twice the total number of cases and controls. For the alternative hypothesis, we used Fisher's noncentral hypergeometric distribution to simulate the number of samples with the pathogenic mutations using the R package *BiasedUrn*[5]. For the null hypothesis, we used the hypergeometric distribution to simulate the number of samples with the pathogenic mutations. Different odds parameters of 40, 60, and 80 were used. In total, there were six different parameter combinations for the number of causal genes and odds parameters. We compared six different methods adjusting for multiple testing. These included two adopted resampling based FDR control methods: RBH_P and RBH_UL. respectively. We also included the BH method[6], and BH_T2 and BH_T3, which first filtered the genes with less than 2 or 3 samples with pathogenic mutations and then applied the BH method to the rest of the genes. We also included a method ADBH.sd recently developed for FDR control of FET[7]. The empirical FDR was the proportion of false positives among detected or 0 if none was detected. The empirical power was the proportion of true positives detected among all true positives. In total, 100 replications were performed, and the final average FDR and power were reported.

##### Type I error and power simulation for LD detection

We simulated six groups of summary counts, with the number of haplotypes being 2*e*4, 2*e*4, 5*e*3, 5*e*3, 3*e*4, 3*e*4, close to the sample size of stratified groups in the nfe population in gnomAD. The AFs of both variants were set to 0.001, 0.005, or 0.01. The odds ratio was set to 1 for evaluating the type I error and 10, 100, 1*e*3, 1*e*4, or 1*e*5 for evaluating the power. The HWE was assumed, and 1,000 replicates were simulated to estimate the type I error and power. The nominal p-value threshold was set to 0.05. We also evaluated the type I error and power when genotypes were observed. We simulated 110,000 haplotypes summed into genotypes per two haplotypes and set the odds ratio to 1, 2, 5, 10, 20, or 50 for evaluation of type I error and power.

##### Simulation of sample-count estimation including independent and correlated variants

To show that identifying high-LD variants can improve sample-count estimation based on summary counts, we simulated seven variants: the first two and the last two were fully correlated and the rest were independent, assuming the HWE. The number of individuals was set to 50,000.

For each simulated data, there were five variants with AF = $p$, including two fully correlated variants, two fully correlated variants with AF = $2p$, and one variant with AF = $p/2$. The parameter $p$ ranged from 0.01 to 0.001, each with three replicates in the simulated data sets. The root mean square error was calculated between the estimated counts and the ground truth. We assumed the frequencies of genotypes was observed in the estimation. We compared CoCoRV with TRAPD on count estimation. The method used in TRAPD overestimated the counts in controls to be conservative in the association test results when using a one-sided FET. For the dominant model, TRAPD added all qualified counts of each variant in a gene; for the double-heterozygous model (called compound heterozygous in Guo *et al.*[8]), the sum of the frequencies of heterozygous genotypes among all variants was squared and then multiplied by the total number of controls; for the recessive model, TRAPD added the counts of homozygous alternate genotypes and the counts from the double-heterozygous model.

**Data processing and comparison between jointly called full genotype–based and summary count–based analysis**

We used the two pediatric cancer cohorts (CNS and ALL) and our constructed in-house controls to compare the concordance between analyses using jointly called full-genotype data and that using separately called summary counts. All sequencing data were remapped using BWA v0.7.12[9] to the reference genome GRCh37-lite. We used GATK v3.7[10] to generate gVCF files for each sample. Three jointly called genotype data sets were generated: 1) full genotype data with all cases and controls jointly called, 2) in-house controls jointly called, and 3) case cohorts jointly called. The variants were normalized, and the multiallelic variants were decomposed into biallelic variants. We followed the GATK3 best practice to apply VQSR on each data set to QC variants. We used the functions *pcair* and *pcrelate* from the R package GENESIS v2.4.0[11] to calculate the PCs and estimated the relatedness from the cases and controls by using the joint genotype calls. After excluding related samples and considering individual sample missingness, there were 8,175 samples in the constructed controls (5,602 from ADSP and 2,573 from the 1,000 Genomes Project). The CNS cohort had 336 cases and the ALL cohort had 958 cases.
